# Kupffer cells control neonatal hepatic metabolism via Igf1 signaling

**DOI:** 10.1101/2025.02.19.638983

**Authors:** Nikola Makdissi, Daria Hirschmann, Aleksej Frolov, Inaam Sado, Fabian Nikolka, Jingyuan Cheng, Nelli Blank-Stein, Maria Francesca Viola, Mohamed Yaghmour, Philipp Arnold, Lorenzo Bonaguro, Matthias Becker, Christoph Thiele, Felix Meissner, Karsten Hiller, Marc D. Beyer, Elvira Mass

## Abstract

During perinatal development, liver metabolism is tightly regulated to ensure energy supply for the newborn. Before birth, glycogen is stored in hepatocytes and later metabolized to glucose, meeting the energy demands of the neonate. Shortly after birth, lipogenesis begins, driven by the transcriptional activation of enzymes involved in fatty acid oxidation. These processes are thought to be largely regulated by systemic insulin and glucagon levels. However, the role of liver-derived local factors in neonatal hepatocyte metabolism remains unexplored. Kupffer cells (KCs), the liver’s resident macrophages, colonize the fetal liver early in embryogenesis and support liver metabolism in adulthood. Yet, whether KCs influence neonatal hepatocyte metabolism is unknown. Here, using conditional knockout mouse models targeting macrophages, we demonstrate that yolk sac-derived KCs play a critical role in hepatocyte glycogen storage and function by regulating the tricarboxylic acid (TCA) cycle - a role that monocyte-derived KC-like cells cannot substitute. Newborn pups lacking yolk sac-derived KCs mobilize glycogen more rapidly, a process regulated by insulin-like growth factor 1 (Igf1) production. Our findings reveal that macrophages are a major source of Igf1 at birth and that local Igf1 production by KCs is essential for balanced hepatocyte metabolism.

## Introduction

The liver is a highly versatile organ that shifts from being the main site of hematopoiesis during embryogenesis to one of the major metabolically active organs in the body after birth (Nakagaki et al., 2018). In neonates, the liver undergoes a metabolic adaptation to support postnatal energy demands (Gong et al., 2020; Li et al., 2023). Before birth, starting at embryonic day (E)17, glycogen is stored in fetal hepatocytes (Tye and Burton, 1980). These stores are mobilized immediately after birth to supply glucose to maintain blood glucose levels during the initial hours prior to milk intake. Within just a few hours, this glycogenolysis phase shifts to active fatty acid oxidation (FAO) and engagement of the TCA cycle, which, together with the supply of amino acids and essential co-factors, fuel gluconeogenesis to maintain glucose homeostasis as milk-derived dietary lipids become the primary energy source. Concurrently, neonatal hepatocytes shift away from glycolysis toward oxidative phosphorylation, maximizing ATP production to meet the high metabolic requirements of the newborn (Li et al., 2023). This metabolic switch ensures a steady glucose supply essential for neonatal tissue homeostasis.

Neonatal liver metabolism is controlled by a combination of hormonal, genetic, and environmental factors. The hormones insulin and glucagon play central roles in inducing glycogen breakdown and gluconeogenesis. After birth, glucagon levels rise sharply, stimulating glycogenolysis and gluconeogenesis, while insulin levels remain low until lactation begins. Lactation provides triglycerides (TG), which are broken down into fatty acids and become the primary energy source for ATP production through FAO and the TCA cycle. Simultaneously, hepatocytes undergo maturation marked by the upregulation of key transcription factors and the expression of metabolically essential genes, which together support increased metabolic activity (Gong et al., 2020; Yang et al., 2023).

Before birth, the liver upregulates the expression of the insulin receptor (Insr) (Gong et al., 2020) to respond to increasing insulin levels and the demand for FAO. In addition to insulin, insulin-like growth factor 1 (Igf1) is a natural ligand of the Insr. Igf1 is an important factor regulating growth and metabolism and is closely linked to insulin signaling pathways (Kineman et al., 2018, 2024). In the adult organism, hepatocytes are the primary source of circulating Igf1 (Kineman et al., 2018, 2024). However, a single-cell atlas of perinatal mouse development identified macrophages, and specifically liver-resident Kupffer cells (KCs), as major producers of Igf1 in the liver (Qiu et al., 2024). While mature hepatocytes do not express significant levels of the Igf1 receptor (Igf1r), its expression is detectable during the early stages of hepatocyte development (Waraky et al., 2016). This suggests that during development, KC-derived Igf1 may influence hepatocyte maturation or metabolic function through Insr/Igf1 signaling pathways. Recent studies further support this hypothesis by demonstrating that macrophage-derived Igf1 in the brain and gut plays a critical role in tissue development (Yan et al., 2022; Rusin et al., 2024).

KCs originate from yolk sac-derived erythro-myeloid progenitors (EMPs) (Gomez Perdiguero et al., 2015). These EMPs differentiate into pre-macrophages (pMacs), which migrate to the liver and establish the early KC population as early as E10.25 (Mass et al., 2016) - preceding the differentiation of hepatoblasts into hepatocytes. We have recently shown that KCs play a vital role in supporting fetal liver hematopoiesis, which peaks at E13.5-E14.5 (Kayvanjoo et al., 2024), underscoring the importance of macrophages in organ development and function (Mass et al., 2023). During adulthood, KCs are also known to regulate liver metabolism through insulin-like growth factor binding protein 7 (Igfbp7) (Morgantini et al., 2019). However, their role in neonatal hepatocyte maturation and metabolism remains unexplored.

In this study, we investigated the role of KCs and KC-derived factors in the development and function of neonatal hepatocytes. Using two KC depletion models and conditional Igf1 knockout mice, we found that KCs are important for proper metabolic function of neonatal hepatocytes. KC depletion accelerated hepatocyte maturation and disrupted metabolic regulation at the transcriptional level. This dysregulation was reflected functionally by reduced TCA cycle activity, diminished insulin responsiveness, resulting in reduced glycogen storage in hepatocytes. Notably, these functions were not restored by KC-like cells that repopulated the empty KC niche. Additionally, depletion of KC-derived Igf1 also led to a decreased glycogen content. Together, these findings reveal that yolk sac-derived KCs and their production of Igf1 are crucial for neonatal hepatocyte development and function.

## Results

### Characterization of Kupffer cell depletion models

To address the role of KCs in neonatal hepatocyte metabolism, we employed two different mouse models that lead to the depletion of tissue-resident macrophages: *Tnfrsf11a^Cre^; Spi1^fl/fl^* and *Tnfrsf11a^Cre^; Csf1r^fl/fl^*(Figure 1A). Using the *Tnfrsf11a^Cre^*, which targets pMacs and therefore all macrophages during development (Mass et al., 2016), we depleted either the transcription factor Spi1 or the surface receptor Csf1r, which are required for macrophage development and survival (Cox et al., 2021; Kayvanjoo et al., 2024; Jacome-Galarza et al., 2019). While we could confirm a complete depletion of F4/80^high^CD11b^int^ KCs at E14.5 in *Tnfrsf11a^Cre^; Spi1^fl/fl^* livers (*KO^Spi1^*) compared to littermates (*WT^Spi1^*) (Kayvanjoo et al., 2024) (Figure 1B, C, Figure S1), the peak of depletion in *Tnfrsf11a^Cre^; Csf1r^fl/fl^* embryos (*KO^Csf1r^*) was detected at E12.5 (Figure 1D, E). At birth (postnatal day (P)0), the livers of *KO^Spi1^* and *KO^Csf1r^* animals were repopulated by F4/80^high^CD11b^int^ cells (Figure 1 F-H), with *KO^Csf1r^* pups showing even significantly increased macrophage numbers (Figure 1H), a phenomenon that has been previously observed after the depletion of tissue-resident macrophages via blockade of Csf1r (Elmore et al., 2018).

**Figure 1.**
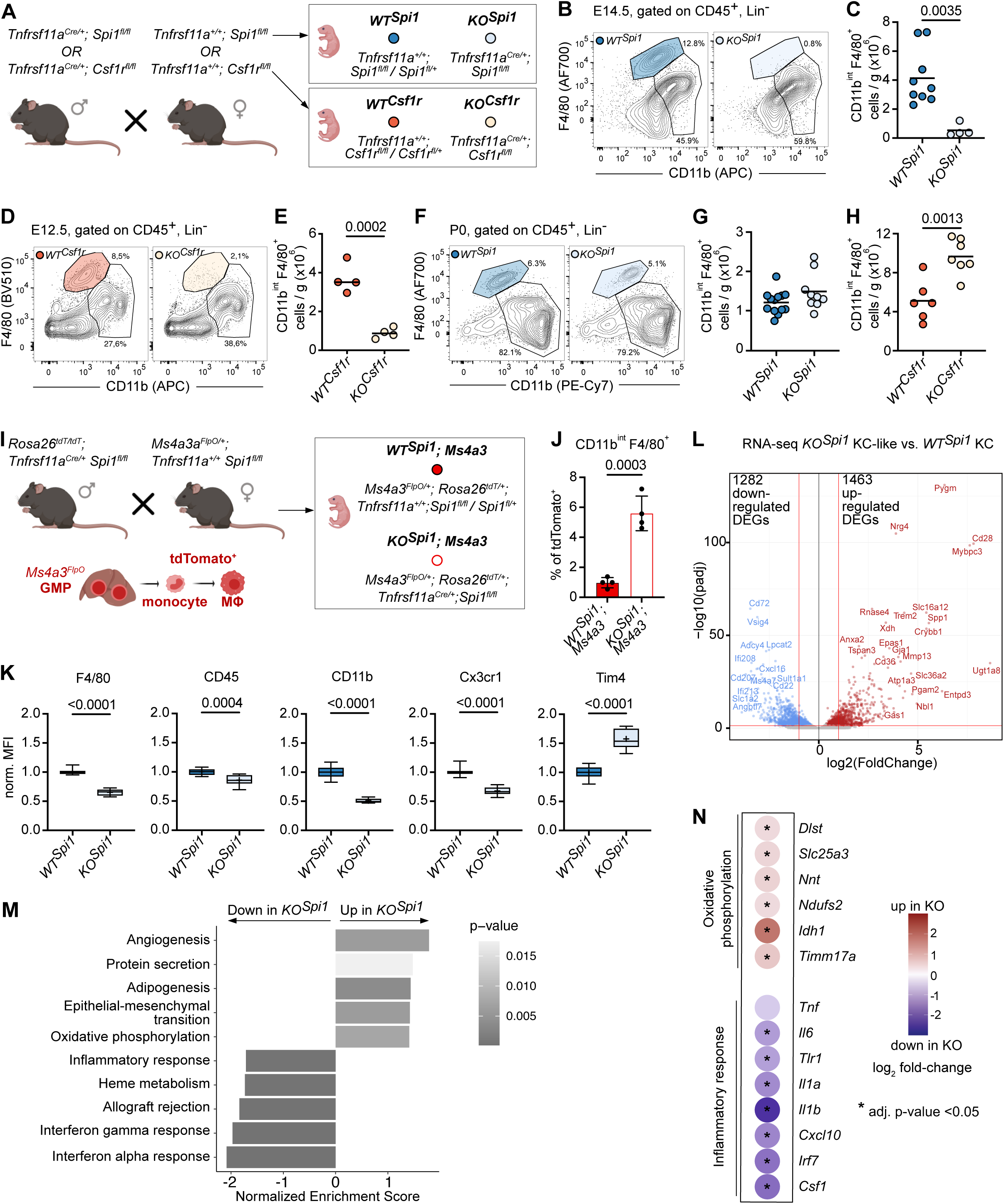
Characterization of the *KO^Spi1^* and *KO^Csf1r^* mouse models. **(A)** Breeding scheme to produce *KO^Spi1^* and *KO^Csf1r^* mice and littermate controls (*WT^Spi1^* and *WT^Csf1r^*). Created with BioRender.com. **(B)** Representative flow cytometry plots of *WT^Spi1^*and *KO^Spi1^* fetal livers at E14.5 showing efficient depletion of KCs. **(C)** Quantification of total *WT^Spi1^* and *KO^Spi1^* KC numbers at E14.5. Circles represent individual mice. n = 4-9 per genotype. Unpaired Student’s t-test. **(D)** Representative flow cytometry plots of *WT^Csf1r^* and *KO^Csf1r^*fetal livers at E12.5 showing efficient depletion of KCs. **(E)** Quantification of total *WT^Csf1r^* and *KO^Csf1r^* KC numbers at E12.5. Circles represent individual mice. n=4 per genotype. Unpaired Student’s t-test. **(F)** Representative flow cytometry plots of *WT^Spi1^* and *KO^Spi1^* livers at P0 showing repopulation of the empty KC niche. **(G)** Quantification of total *WT^Spi1^* and *KO^Spi1^*KC numbers at P0. Circles represent individual mice. n = 9-13 per genotype. **(H)** Quantification of total *WT^Csf1r^* and *KO^Csf1r^* KC numbers at P0. Circles represent individual mice. n = 9-13 per genotype. Unpaired Student’s t-test. **(I)** Breeding scheme to produce *KO^Spi1^* mice and littermate controls in combination with the granulocyte-monocyte progenitor (GMP) fate-mapping model *Ms4a3^FlpO^*. Labelled GMPs differentiate to monocytes and subsequently to macrophages (M<λ). **(J)** Quantification of fate-mapped KCs and KC-like cells in *WT^Spi1^* and *KO^Spi1^*mice at P0. Circles represent individual mice. n = 4 per genotype from 2 litters. Unpaired Student’s t-test. **(K)** Normalized expression of surface receptor markers on KCs and KC-like cells from *WT^Spi1^* and *KO^Spi1^* mice at P0. n = 9-13 per genotype. Boxplot with 5-95 percentile. Cross indicates the mean, and the line the median. Mann-Whitney test**. (L)** Volcano plot showing differentially expressed genes (DEGs) comparing KCs from *WT^Spi1^*and KC-like cells from *KO^Spi1^* mice at P0. n = 5 per genotype. log_2_FoldChange=1.3, p-adjust=0.05. **(M)** Gene Ontology enrichment analysis of differentially up- and down-regulated genes comparing KCs from *WT^Spi1^* and *KO^Spi1^* mice at P0. **(N)** Selected genes from the terms ‘Oxidative phosphorylation’ and ‘Inflammatory response’ from (L). Dotplot indicates up-(red) and down-regulated (blue) genes when comparing *KO^Spi1^*to *WT^Spi1^*. Wald test for pairwise comparisons. * adjusted p-value <0.05.

To investigate whether the depleted KC niche would be repopulated by EMP-derived cells or by an alternative source, such as hematopoietic stem cell (HSC)-derived monocytes, we utilized a genetic labeling approach. In this context, it is important to note that while KCs primarily arise from yolk sac-derived EMPs, HSC-derived monocytes can also serve as a source for macrophages under certain conditions, including injury or inflammation (Mass et al., 2023; Schultze et al., 2019). To distinguish between these two potential sources, we crossed the *KO^Spi1^*model to the *Ms4a3^FlpO^; Rosa26^FLF-tdTomato^* model (Figure 1I), which specifically labels cells originating from the definitive HSC wave (Huang et al., 2023; Liu et al., 2019). This allowed us to trace the contribution of HSC-derived monocytes in the repopulation process. While the number of tdTomato-positive cells increased significantly from ∼1% to 5-7%, a large fraction remained tdTomato-negative (Figure 1J). This indicates that EMP-derived monocytes (Gomez Perdiguero et al., 2015) or other macrophage precursors residing in the fetal liver can readily differentiate into KC-like cells.

Although these KC-like cells expressed high levels of F4/80 (see gating in Figure 1F), the relative expression of F4/80 as well as other macrophage markers, such as CD45, CD11b and Cx3cr1 were significantly reduced (Figure 1K). In contrast, Tim4 expression was increased (Figure 1K). Transcriptome analysis of sorted F4/80^high^CD11b^int^ KCs and KC-like cells from of P0 *KO^Spi1^* and *WT^Spi1^*livers, respectively, showed that the freshly differentiated KC-like cells had an altered transcriptional profile (Figure S2A, B), resulting in 1282 downregulated and 1463 upregulated differentially expressed genes (DEGs) (Figure 1L, Table 1). Gene set enrichment analysis indicated that genes falling into terms such as protein secretion, angiogenesis, and oxidative phosphorylation were enriched in *KO^Spi1^* KC-like cells while genes belonging to inflammatory response, heme metabolism, and interferon alpha and gamma responses were downregulated (Figure 1M, Table 2). Upregulated DEGs falling into the term ‘oxidative phosphorylation’ included *Slc25a3*, *Ndufs2*, and *Idh1*, whereas *KO^Spi1^* KC-like cells downregulated genes such as *Tnf, Il6, Il1a, Il1b*, and *Cxcl10* falling into the term ‘inflammatory response’ (Figure 1N). These findings, along with the altered surface expression of macrophage markers, demonstrate that the newly formed KC-like cells possess a distinct molecular composition compared *to bona fide* EMP-derived KCs. In summary, the *KO^Spi1^* and *KO^Csf1r^* models represent two KC-depletion models that allow to study the role of macrophage ontogeny in tissue development and homeostasis.

### KC depletion leads to decreased glycogen storage at birth

After establishing the KC-depletion models, we investigated whether the replacement of long-lived KCs by KC-like cells could affect neonatal hepatic metabolism. The liver plays a critical role in energy conversion, maintaining blood glucose homeostasis in neonates by storing glycogen during embryogenesis, which is subsequently metabolized through extensive glycogenolysis after birth (Li et al., 2023). Therefore, we assessed glycogen levels in P0 *WT^Spi1^*and *KO^Spi1^* livers using PAS staining (Figure 2A). Quantification of the staining indicated that *KO^Spi1^* livers stored less glycogen than their littermate controls (Figure 2B, C). For a more quantitative analysis, we used a colorimetric assay, which corroborated our histological results, showing that *KO^Spi1^* livers had approximately half of the glycogen concentration of *WT^Spi1^* livers (Figure 2D). Moreover, transmission electron microscopy images clearly demonstrated a reduction in glycogen content in *KO^Spi1^*compared to *WT^Spi1^*, where the glycogen appears more abundant and densely packed (Figure 2E). We further confirmed a significantly reduced glycogen content in the *KO^Csf1r^*livers compared to *WT^Csf1r^* using colorimetric assay (Figure 2F). Collectively, our findings suggest that the replacement of depleted KCs by KC-like cells leads to an altered hepatocyte metabolism at birth, characterized by significant changes in glycogen storage.

**Figure 2.**
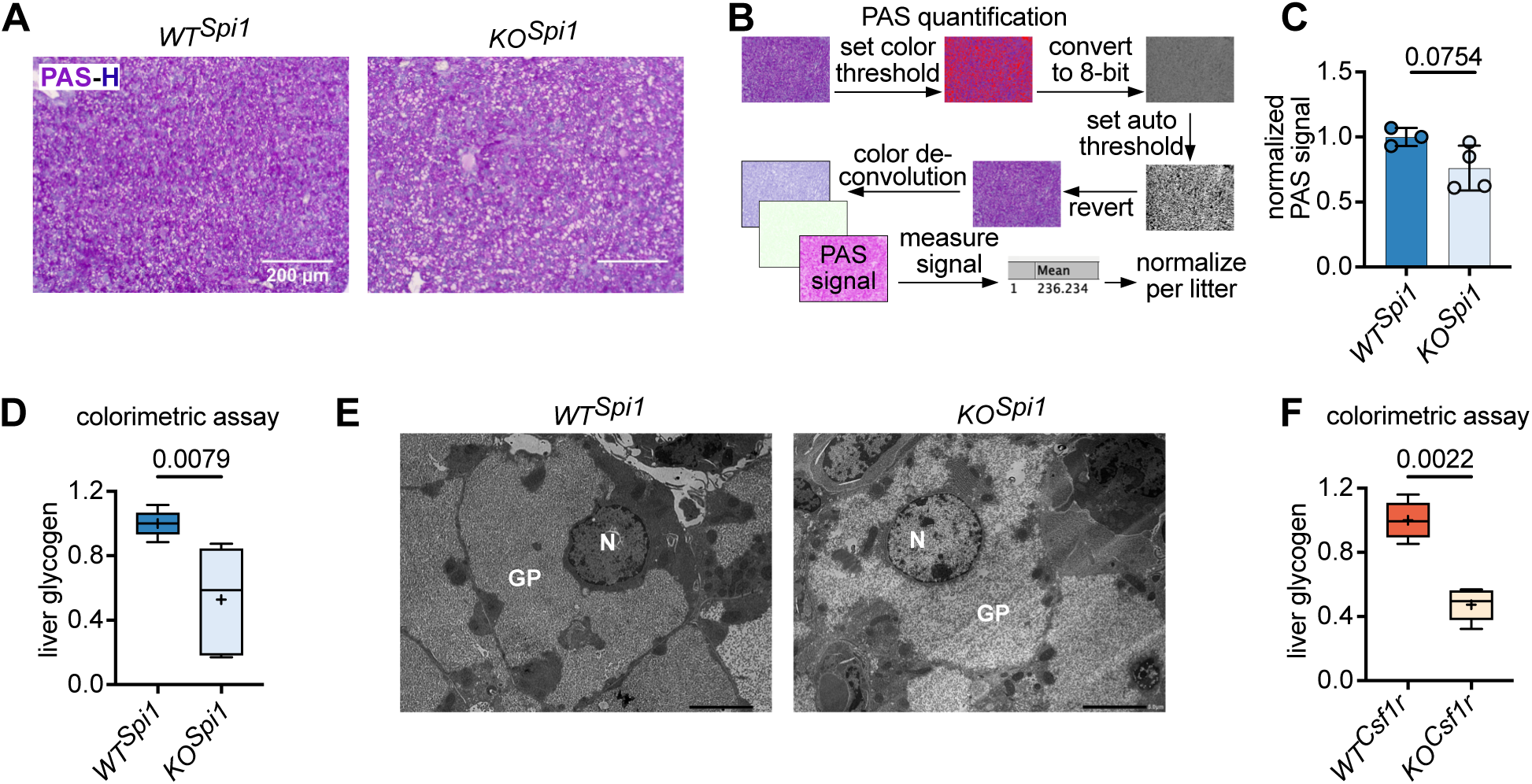
*KO^Spi1^*and *KO^Csf1r^* hepatocytes show less glycogen storage at birth. **(A)** Representative PAS (purple)-Hematoxylin (H, blue) staining of *WT^Spi1^* and *KO^Spi1^* livers at P0. Scale bar = 200 µm. **(B)** Scheme indicating how PAS staining intensity was quantified using Fiji. **(C)** Normalized PAS signal intensity of *WT^Spi1^*and *KO^Spi1^* livers at P0. Circles represent individual mice. n=3-4 per genotype from 2 litters. Paired Student’s t-test. **(D)** Glycogen levels measured on whole liver lysates of *WT^Spi1^* and *KO^Spi1^* P0 livers. n=5 per genotype from 3 litters. Values were normalized per litter. Boxplot with 5-95 percentile. Cross indicates the mean, and the line the median. Mann-Whitney test**. (E)** Representative transmission electron micrograph from *WT^Spi1^* and *KO^Spi1^* livers at P0. n = 3 per genotype. GP: glycogen particle; N: nucleus. Scale bar = 5 µm. **(F)** Glycogen levels measured on whole liver lysates of *WT^Csf1r^*and *KO^Csf1r^* at P0. n = 6 per genotype from 3 litters. Values were normalized per litter. Boxplot with 5-95 percentile. Cross indicates the mean, and the line the median. Mann-Whitney test.

### Hepatocytes undergo a metabolic shift after Kupffer cell depletion

To understand whether hepatocytes are altered functionally due to the lack of KCs, we first performed single-cell RNA-sequencing (scRNA-seq) analysis of livers from P0 *WT^Spi1^* and *KO^Spi1^* (Figure S2C, D). We identified hepatocytes based on lineage markers (Figure S2D). After identifying all DEGs between *WT^Spi1^* and *KO^Spi1^* we ranked these genes based on average log_2_ fold-change and used the top 100 up- or downregulated genes for Gene ontology (GO) enrichment analysis (Table 3). The results of this analysis showed a significant upregulation of ‘lipid transport’, ‘ATP metabolic process’, ‘pyruvate metabolic process’, and ‘response to insulin’, and downregulation of ‘fatty acid metabolic process’ and ‘gluconeogenesis’ terms in *KO^Spi1^* hepatocytes (Figure 3A).

**Figure 3.**
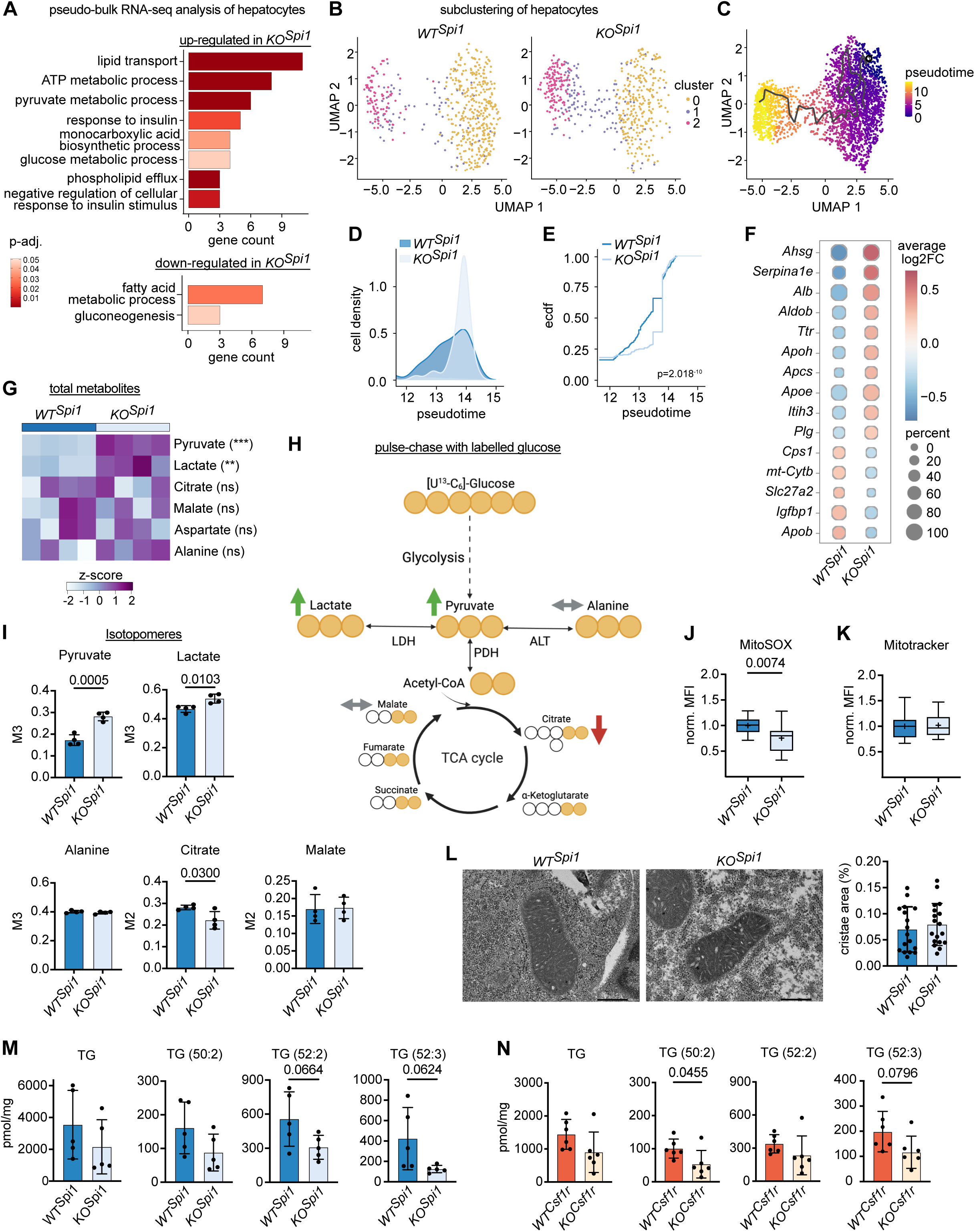
*KO^Spi1^*hepatocytes exhibit an altered metabolism following Kupffer cell depletion. **(A)** Pseudo-bulk RNA-Seq analysis of hepatocytes showing gene ontogeny enrichment for up- and downregulated pathways in *KO^Spi^* at P0. **(B)** Uniform Manifold Approximation and Projection (UMAP) analysis of *WT^Spi^* and *KO^Spi^* hepatocytes from the scRNA-Seq dataset. **(C)** Pseudotime trajectory analysis of the hepatocytes cluster using Monocle3. **(D)** Density distribution analysis of hepatocytes within the pseudotime interval 12-15. **(E)** Empirical cumulative distribution function (ecdf) analysis of hepatocytes within the pseudotime interval 12-15. **(F)** Pseudo-bulk analysis of DEGs in hepatocyte cluster 2. Dotplot shows log_2_ fold-change (color) and percentage of cells expressing each gene (dot size). **(G)** Normalized total metabolite abundance in of *WT^Spi^* and *KO^Spi^* livers following ^13^C-glucose tracing at P0. n = 4 per genotype from 2 litters. Unpaired Student t-test. *p<0.05, **p<0.01, ***p<0.001. **(H)** Schematic representation of the possible metabolite labeling patterns due to the incorporation of the ^13^C-glucose tracer. Created with BioRender.com. **(I)** Fractional enrichment of labeled metabolites following ^13^C-glucose tracing at P0. n= 4 per genotype from 2 litters. Barplot presented as mean ± SD. Unpaired Student t-test. **(J, K)** Normalized median fluorescent intensity (MFI) of MitoSOX Red **(J)** and Mitotracker Green **(K)** in hepatocytes at P0. n = 13-14 per genotype from 5 litters. Boxplot with 5-95 percentile. Cross indicates the mean, and the line the median. Unpaired Student t-test. **(L)** Representative transmission electron micrographs of mitochondria of *WT^Spi1^* and *KO^Spi1^* livers (left), along with the quantified cristae area coverage (right). n = 17-18 mitochondria. Scale bar = 500 nm. **(M, N)** The abundance of total triacylglycerol (TG) and its subspecies in *WT^Spi^* and *KO^Spi^* **(M)** and *WT^Csf1r^* and *KO^Csf1r^* **(N)** livers. n = 5-6 per genotype from more than 3 litters per mouse model. Barplot presented as mean ± SD. Unpaired Student t-test.

Given the dysregulation of metabolic pathways at the transcriptional level and the accelerated glycogen degradation, we aimed to determine whether the absence of KCs affects hepatocyte maturation in *KO^Spi1^* animals. To this end, we subclustered all hepatocytes, which resulted in three distinct clusters (Figure 3B). We examined the developmental trajectory of hepatocytes using trajectory inference with Monocle3. To rank cells on a pseudotemporal scale we performed pseudotime analysis with a cell from cluster 0 as root node, because cells in this cluster expressed the lowest levels of hepatocyte maturation markers, such as *Alb*, *Aldob*, *Ahsg*, and *Ttr* (Gong et al., 2020). Thus, in this analysis, cluster 0 represents the immature state, and progresses to cluster 2, which represents the most mature state (Figure 3C). As the distribution of cells among cluster 2 was altered in *KO^Spi1^* hepatocytes (Figure 3B), we ranked genes expressed by cluster 2 from *WT^Spi1^* and *KO^Spi1^* hepatocytes based on average log_2_ fold-change and used the top 100 up- or downregulated genes for GO enrichment analysis. This analysis revealed that distinct genes falling into the same terms were either upregulated in *WT^Spi1^* or in *KO^Spi1^* hepatocytes (Figure S2E). Common terms included ‘fatty acid metabolic process’ and ‘gluconeogenesis’ with genes such as *Lpin1* and *Lpin2*, which are important for TG biosynthesis, showing a reduced expression in *KO^Spi1^*hepatocytes, while other genes related to fatty acid metabolic processes, such as *Apoa1* and *Lpl,* were expressed at higher levels in *KO^Spi1^*hepatocytes (Figure S2E). These data suggest that *KO^Spi1^* hepatocytes, despite belonging to the most mature cluster, shifted their metabolic functionality compared to *WT^Spi1^* hepatocytes. To further address this transcriptional shift, we performed a density distribution (Figure 3D) and an empirical cumulative distribution analysis (Figure 3E) focusing on the pseudotime interval 12-15. These analyses revealed that *KO^Spi1^* hepatocytes progressed more rapidly toward the mature state. We corroborated this by performing a pseudo-bulk analysis of DEGs in cluster 2: the upregulation of hepatocyte maturation markers, such as *Alb, Aldob, Ahsg*, and *Ttr* (Gong et al., 2020) in *KO^Spi1^* hepatocytes compared to *WT^Spi1^* (Figure 3F) further supported the hypothesis that KC-deficiency during embryogenesis promotes hepatocyte maturation on the transcriptional level.

To determine whether *KO^Spi1^* hepatocytes exhibit increased metabolic activity due to their accelerated transcriptional maturation, we performed 5-hour ^13^C-glucose tracing experiments on freshly isolated livers from *WT^Spi1^* and *KO^Spi1^* mice. Normalized metabolite abundance showed significantly higher levels of pyruvate and lactate in *KO^Spi1^* livers, while alanine, aspartate, citrate and malate remained comparable between conditions (Figure 3G). ^13^C-labelled metabolites also showed increased isotope incorporation into pyruvate and lactate (Figure 3H, I); however, glycolytic acetyl-CoA incorporation into citrate (M2) was significantly lower in *KO^Spi1^* livers, while contribution to malate was similar across genotypes (Figure 3H, I). Collectively, these findings suggest a reduced pyruvate dehydrogenase (PDH) flux and altered carbon flow into the TCA cycle in *KO^Spi1^* livers. Moreover, hepatocytes showed decreased reactive oxygen species production, as measured by MitoSOX staining (Figure 3J). The MitoSOX signal serves as a proxy for mitochondrial respiration measurement and, therefore, indicates reduced TCA cycle activity, possibly due to the decreased PDH flux. Of note, measurement of MitoTracker green fluorescence (Figure 3K), indicative for mitochondrial quantity, and quantification of the cristae area coverage in mitochondria using transmission electron microcopy (Figure 3L), did not reveal any changes between *WT^Spi1^* and *KO^Spi1^*livers. Additionally, high-dimensional flow cytometry of hepatocytes did not detect significant changes in the abundance of various nutrient transporters (GLUT1, CD36, CD98) and core metabolic enzymes (PKM, G6PD, SDHA, ATP5a, CPT1a, ACC1) (Figure S3A-C). These data confirm that the altered metabolic activity arises from changes in the metabolic response of hepatocytes rather than from impairments in mitochondrial health. In summary, our findings from metabolic tracing experiments indicate that *KO^Spi1^* hepatocytes have enhanced glycolytic activity and increased production of pyruvate and lactate, yet exhibit reduced TCA cycle engagement as evidenced by lower citrate incorporation.

To further explore whether the changes in PDH flux and TCA cycle activity impact lipid metabolism, we performed a comprehensive lipid analysis using mass spectrometry. Overall, no significant changes were found in lipid classes across both KO models (Figure S3D, E). However, we observed a reduction in the most abundant TG species, including TG(50:2), TG(52:2), and TG(52:3), in both *KO^Spi1^* and *KO^Csf1r^* livers (Figure 3M, N), indicating reduced synthesis of these specific lipid types. Collectively, our data demonstrates the requirement of KCs for proper neonatal hepatocyte function, as the presence of KC-like macrophages leads to a metabolic dysregulation, with a preference for lactate production over TCA cycle and lipid synthesis. This shift may also explain the need of increased consumption of glycogen to maintain ATP levels, leading to faster glycogen depletion (see Figure 2).

### KC-derived Igf1 controls glycogen homeostasis in neonatal hepatocytes

To identify KC-dependent pathways potentially driving the metabolic shift and increased glycogen demand in hepatocytes, we used the scRNA-seq data from *WT^Spi1^* and *KO^Spi1^* livers and performed CellChat analysis, focusing on the interaction between KCs and hepatocytes (Figure 4A). The analysis revealed several downregulated pathways in KC-like macrophages from *KO^Spi1^* animals, particularly those involved in regulating cellular metabolism, including the VISFATIN, IGF, and SEMA6 pathways (Kang et al., 2018; Nakanishi et al., 2024; Revollo et al., 2007; Kineman et al., 2024; Fellinger et al., 2023; Kineman et al., 2018). Notably, visfatin (also known as the extracellular form of Nampt) has been shown to regulate insulin secretion (Revollo et al., 2007). To investigate whether KC-like macrophages from *KO^Spi1^* livers interact uniformly with all hepatocyte populations or if specific hepatocyte clusters are more affected, we analyzed ligand-receptor pairs to explore these dynamics (Figure 4B). Our analysis revealed that Nampt-Insr interactions had the highest probability of interaction between *WT^Spi1^* KCs and all three hepatocyte clusters, but were markedly reduced or absent in *KO^Spi1^* livers (Figure 4B). *Igf1-Igf1r* signaling was highest between *WT^Spi1^* KCs and hepatocyte cluster 2 (Hepa2), but showed reduced communication probability between KCs and Hepa2 in *KO^Spi1^* livers. Similar behavior was observed for the ligand-receptor pair *Sema6a-Plxna2* (Figure 4B). Although not listed among the main signaling pathways altered between *WT^Spi1^* and *KO^Spi1^* animals (Figure 4A), we predicted a potential interaction of Tnf and Tnfrsf1a in *WT^Spi1^* livers based on the *Tnf* gene expression in KCs and *Tnfrsf1a* gene expression in all hepatocyte clusters (Figure 4B). This interaction was absent in *KO^Spi1^* livers (Figure 4B). This further supports the hypothesis that neonatal yolk sac-derived KCs play a role in regulating hepatocyte metabolism, as KC-derived Tnf is known to control lipid uptake in hepatocytes (Diehl et al., 2020).

**Figure 4:**
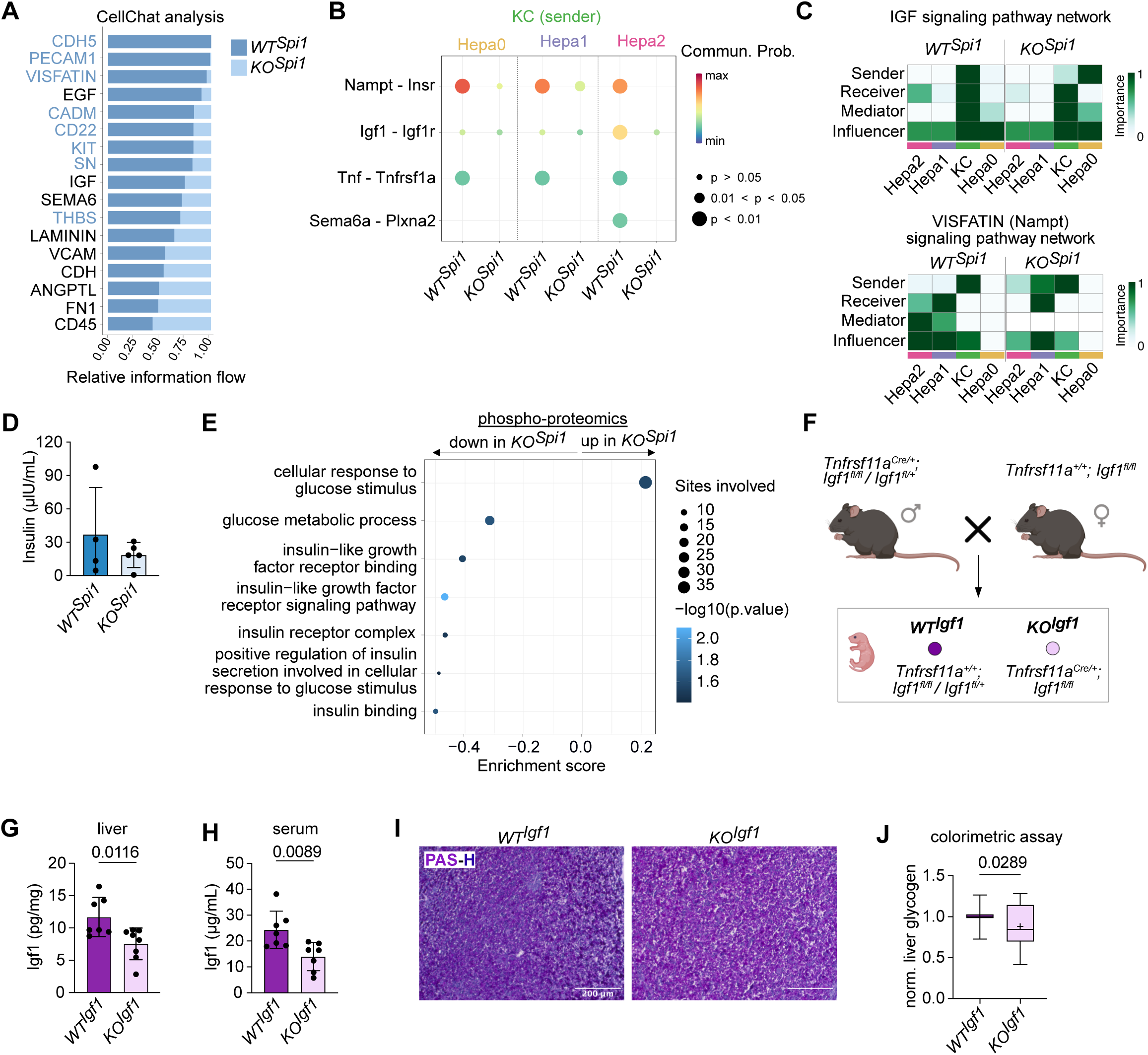
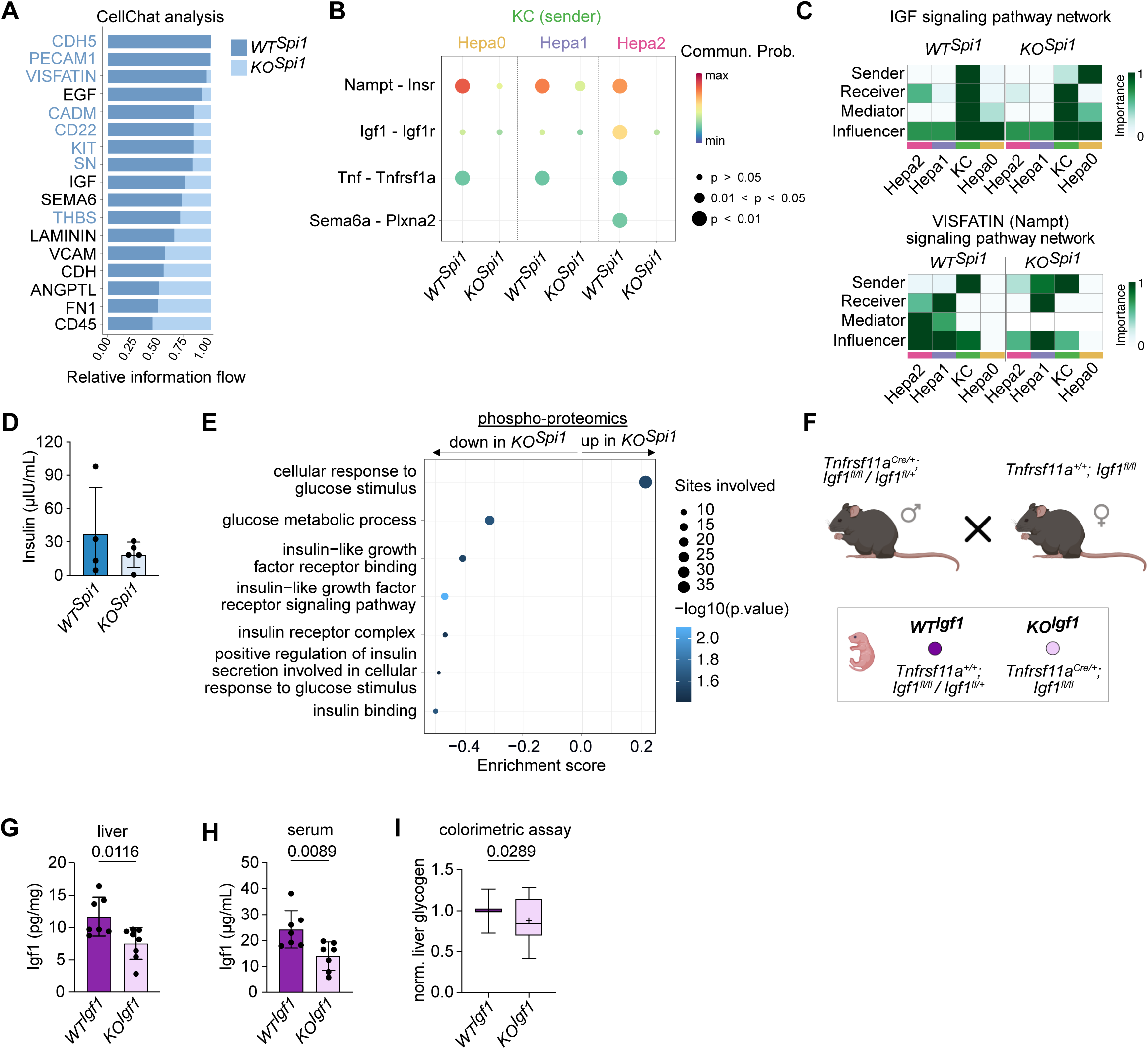
KC-derived Igf1 regulates glycogen homeostasis in neonatal hepatocytes. **(A)** Bar plot showing the relative information flow of between *WT^Spi^* and *KO^Spi^* of inferred cell-cell communication using CellChat. **(B)** Comparison of the significant ligand-receptor pairs between *WT^Spi^* and *KO^Spi^*, which contribute to the signaling from Kupffer cells to the hepatocyte clusters. **(C)** Heatmap shows the relative importance of Kupffer cell and hepatocyte clusters as sender, receiver, mediator and influence, based on the computed four network centrality measures of IGF (upper) and VISFATIN (lower) signaling. **(D)** Serum insulin levels measured by ELISA on *WT^Spi1^* and *KO^Spi1^* at P0. n = 4-5 per genotype from 4 litters. Barplot presented as mean ± SD. Unpaired Student t-test. **(E)** Enrichment analysis of downregulated phosphorylation sites showing the decreased and increased phosphorylation in *KO^Spi1^* liver compared to *WT^Spi1^*. **(F)** Breeding scheme to produce *KO^Igf1^* mice and littermate controls (*WT^Igf1^*). Created with BioRender.com. **(G, H)** Igf1 levels measured by ELISA on whole liver lysate **(G)** and serum **(H)** of *WT^Spi1^* and *KO^Spi1^* at P0. n = 4-5 per genotype from 4 litters. Barplot presented as mean ± SD. unpaired Student t-test. **(I)** Glycogen levels measured on whole liver lysates of *WT^Igf1^* and *KO^Igf1^* at P0. n = 11-16 per genotype from 7 litters. Values were normalized per litter. Boxplot with 5-95 percentile. Cross indicates the mean, and the line the median. Mann-Whitney test.

Next, we analyzed the pathways directly associated with Insr and Igf1r in more detail. Network centrality analysis in *WT^Spi1^* and *KO^Spi1^* KCs and hepatocyte clusters suggested that KCs were the primary senders, mediators, and influencers of the IGF signaling pathway in *WT^Spi1^* animals. In contrast, *KO^Spi1^* KCs exhibited reduced sender activity, with the least mature hepatocyte cluster 0 emerging as a prominent sender (Figure 4C). Analysis of the Visfatin (Nampt) signaling pathway further revealed that the most mature cluster Hepa2 in *KO^Spi1^* livers showed a diminished role in receiving signals via Insr. Additionally, Hepa2 in *KO^Spi1^* livers showed diminished activity as mediator and influencer to its counterpart in *WT^Spi1^*animals (Figure 4C). While Nampt does not directly bind to the Insr, Igf1 interacts with both Insr homodimers and Insr/Igf1r heterodimers. Despite this difference, both Nampt and Igf1 are capable of activating downstream Insr signaling pathways, thereby regulating cellular metabolism (Saddi-Rosa et al., 2010; Kineman et al., 2018). Together, our transcriptional data provides evidence that metabolic function of hepatocytes may be regulated by KCs, which either directly or indirectly modulate the Insr signaling cascade.

Neonatal hepatocytes can sense insulin via the Insr, which promotes glucose uptake (Nevado et al., 2006). Thus, we tested whether a disbalance of systemic insulin levels may drive the metabolic shift seen in *KO^Spi1^* hepatocytes. However, plasma insulin levels did not differ between *KO^Spi1^* and *WT^Spi1^* animals at P0 (Figure 4D). Next, we employed a quantitative phospho-proteomics approach and compared whole livers of *WT^Spi1^*and *KO^Spi1^* littermates at P0. To complement our findings from the scRNA-seq data, we specifically focused on glucose- and insulin-related signaling pathways. Enrichment analysis of downregulated phosphorylation sites comparing *KO^Spi1^* and *WT^Spi1^* livers (Table 4) showed that sites involved in ‘glucose metabolic process’, ‘Igf1 binding’, ‘Igf1 signaling pathway’, and ‘insulin binding’ were less phosphorylated. Conversely, sites involved in ‘cellular response to glucose stimulus’ showed an increased phosphorylation in *KO^Spi1^* livers when compared to *WT^Spi1^* (Figure 4E). Notably, downregulated phosphorylation sites in *KO^Spi1^* livers included key sites on insulin receptor substrate 1 (Irs1) - S307, S318, and S526 (Table 4) - defined as direct downstream activation sites of the Insr (Humphrey et al., 2013; Müssig et al., 2005; Rui et al., 2001; Parker et al., 2015). Additionally, the downregulated GO term ‘insulin receptor complex’ included the site S1340 on the Insr, which is phosphorylated upon insulin stimulation (Parker et al., 2015). In summary, our findings suggest that KC-like macrophages in *KO^Spi1^*livers are unable to engage the signaling pathway downstream of Igf1 and Insr in neonatal hepatocytes as efficiently as *bona fide* KCs, potentially contributing to the observed depletion of hepatic glycogen stores.

Previous studies have shown that macrophages, especially during the early postnatal period, are major producers of Igf1 (Pridans et al., 2018; Yan et al., 2022). To address whether KC-derived Igf1 directly controls glycogenolysis, we generated the macrophage-specific Igf1 knockout mouse model *Tnfrsf11a^Cre^; Igf1^fl/fl^* (Figure 4F). We first confirmed that macrophages were the primary producers of Igf1 at P0. Indeed, conditional knockout mice (*KO^Igf1^*) exhibited significantly lower Igf1 levels in both the serum and liver compared to littermate controls (*WT^Igf1^*, Figure 4G, H). This reduction was accompanied by lower glycogen levels in the liver at P0, as demonstrated via quantitative colorimetric assay (Figure 4I). In summary, these findings suggest that KC-derived Igf1 plays a crucial role in regulating neonatal glycogen metabolism, with the loss of Igf1 leading to a marked reduction in hepatic glycogen stores, highlighting the importance of KC-hepatocyte cross-talk in maintaining metabolic homeostasis during the early postnatal period.

## Discussion

Here, we show a requirement for KC-derived Igf1 in the metabolic homeostasis of the neonatal liver. Depletion of KCs during embryogenesis using two different models (*KO^Spi1^* and *KO^Csf1r^*) led to a replenishment of the empty niche by KC-like macrophages. Despite the presence of macrophages in both models at birth, we observed a marked reduction of glycogen storage in livers from *KO^Spi1^* and *KO^Csf1r^* mice, indicating that hepatocyte function depends on the unperturbed development of EMP/pMac-derived macrophages. Using the Ms4a3 promoter-based fate-mapping in combination with the *KO^Spi1^* model, we confirmed an increased influx of HSC-derived monocytes into the KC niche. However, most of the KC-like macrophages did not stem from definitive HSCs. Recent studies suggest long term (LT)-HSC-independent hematopoiesis in the fetal liver (Kobayashi et al., 2023; Yokomizo et al., 2022), which may partially account for the unlabeled cells in our model as Ms4a3 is only labelling GMPs stemming from definitive LT-HSCs (Liu et al., 2019). Alternatively, EMP-derived monocytes, which are readily found in the fetal liver (Gomez Perdiguero et al., 2015), and which remain Ms4a3-negative, may be the source of new macrophages in the liver. Thus, our combined KO/fate-mapping approach highlights the so far elusive *in vivo* potential of HSC-independent progenitors generating macrophages and the need to develop novel mouse models that can discriminate EMP-from HSC-derived monocytes.

KCs are among the first tissue-specific macrophages that can be detected in the developing embryo as early as E10.25 (Mass et al., 2016). Thus, hepatocytes, which begin differentiating at E13.5 (Yang et al., 2017) and start glycogen storage at E17 (Tye and Burton, 1980), are continuously interacting with KCs, which only begin to migrate to the sinusoids at one week of age (Araujo David et al., 2024). This direct cell communication positions KCs as critical influencers of hepatocyte maturation and function. Using a combination of scRNA-seq, phospho-proteomics, and a conditional knockout of Igf1 in macrophages (*KO^Igf1^*), we identify KCs as active regulators of hepatocyte function. Neonatal KCs produce Igf1, which supports TCA activity and glucose metabolism in hepatocytes before hepatocytes themselves begin producing Igf1. It remains to be determined whether Igf1 exerts its effects on perinatal hepatocyte metabolism via monomeric Igf1r, Insr homodimers, or their heterodimers. The role of macrophage-derived Igf1 in tissue development and function, also observed in the brain and gut (Yan et al., 2022; Rusin et al., 2024), underscores the specialized supportive functions of yolk sac-derived macrophages in their surrounding niche, particularly before growth factor production by other tissue cells commences.

Notably, this study shows that KC-like cells derived from sources other than the yolk sac are unable to perform the same functions. We have previously proposed the concept of developmental programming of macrophages, suggesting that their co-development with niche cells, ontogeny, and longevity are critical factors controlling their effector functions and, consequently, organ development and function (Mass and Gentek, 2021; Mass et al., 2023; Viola et al., 2024). Our findings support the necessity for developmental programming of KCs, which enables them to fulfil their core function in the neonatal liver, namely the regulation of hepatocyte metabolism.

## Methods

### Mice

All investigations involving mice have been locally approved, and performed procedures have followed the guidelines from Directive 2010/63/EU of the European Parliament on protecting animals used for scientific purposes. The experiments were carried out following the German law of animal protection and met the criteria defined by the local institutional animal care committee (Landesamt für Natur, Umwelt und Verbraucherschutz (LANUV), North396 Rhine-Westphalia, Az *81-02.04.2018.A056 & Az 81-02.04.2023.A144*). The mice were house under specific pathogen-free conditions, a 12-h light/dark cycle, with food and water available *ad libtium*. Timed matings were performed overnight females were plug checked the next morning, and pregnant females were monitored regularly. All mice were maintained on a C57BL/6JRCC background. To generate mice lacking macrophages during embryonic development *Tnfrsf11a^Cre^; Spi1^fl/fl^* mice *Tnfrsf11a^Cre^; Spi1^fl/+^* males were crossed with *Spi1^fl/f^* females (a similar scheme was employed for *Tnfrsf11a^Cre^; Csf1r^fl/fl^*mice). For fate-mapping incoming myeloid cells *Tnfrsf11a^Cre^; Spi1^fl/fl^* mice were crossed to *Ms4a3^FlpO^; Rosa26^FLF-tdTomato/FLF-tdTomato^* mice (*Rosa26^FLF-tdTomato^* mice were originally obtained from JAX stock #032864). For prenatal time points pregnant females were sacrificed by cervical dislocation and the embryos were decapitated directly after C-section. All postnatal mice were separated from their mothers and decapitated before organ collection. All mice were genotyped referring to protocols and primers provided by JAX or donating researchers.

### Preparation of single-cell suspension for flow cytometry

Pre- and postnatal mice were sacrificed as described, and the liver was removed and weighed. For E12.5 and E14.5, the whole liver was processed, and for P0 half was used for further steps. The tissue was cut into small pieces (or dissociated with a pipette for E12.5-E14.5), and incubated with 500 µl of a digestion mix (80 U/ml DNase I (Sigma-Aldrich, Cat# DN25) and 1 mg/mL collagenase D (Roche, Cat# 11088858001) in FACS buffer (0.5 % BSA, 2 mmol EDTA in PBS)) for 30 min at 37 °C. All following steps were performed on ice. The suspension was further passed through a 70 µm filter, and 5 ml cold FACS buffer was added. The samples were centrifuged at 400 g for 5 min at 4°C and the supernatant was discarded. Red blood cell lysis was performed only for the P0 samples by dissolving the cells gently in 1 ml of cold RBC lysis buffer (155 mM NH4Cl, 12 mM NaHCO3, and 0.1 mM EDTA) and 5 min incubation. After adding 5 ml cold FACS buffer, the samples were again centrifuged (400 g, 5 min, 4 °C). The supernatant was discarded, 50 μl of Fc-blocking buffer (anti-CD16/32 and 2% rat serum (liver) in FACS buffer) was added and the cells were gently resuspended. After 10 min incubation, all samples were filled up to 250 μl each and counted using a Guava cell counting device. After pelleting the cells were stained for 30 min (both primary and secondary antibody steps). The complete list of antibodies is supplied in Table 5. Samples were acquired with FACSymphony^TM^ A5 (BD Biosciences) and analyzed in FlowJo^TM^ Software.

### Mitochondrial stainings

To evaluate mitochondrial status, liver cells were stained using MitoTracker® Green FM (Thermo Fisher Scientific, M7514) and MitoSOX™ Red (Thermo Fisher Scientific, M36008). Following cell counting, 5 × 10⁵ cells were resuspended in FACS buffer and transferred into a 96-well plate. For stained samples, the volume was adjusted to 50 μL with FACS buffer, while unstained controls were adjusted to 100 μL. Next, 50 μL of prewarmed (37 °C) MitoTracker Green (0.25 μM) and MitoSOX Red (1 μM) solution in HBSS was added to the stained samples. The samples were incubated for 30 minutes at 37 °C, after which 100 μL of FACS buffer was added to all wells. The plate was centrifuged at 400 × g for 5 minutes at 4 °C, and the supernatant was carefully discarded. The cells were then incubated with the respective antibody mix (Table 5) for 30 minutes at 4 °C. Following staining, the samples were immediately acquired on the flow cytometer.

### Metabolic flow cytometry assay

Single-cell suspensions of P0 livers were blocked with anti-CD16/32 (1%) and rat serum (2%) in FACS buffer, followed by surface marker staining for 30 min at 4 °C. Cells were washed and fixed in 1% PFA for 5 minutes, then permeabilised for 15 min with PBS supplemented with 0.4% Triton^TM^ X-100 (Sigma-Aldrich, Cat# X100). Intracellular staining was performed for 1 h, 4 °C. Intracellular antibodies were conjugated in-house using lightning-link conjugation kits (Abcam) as previously described (Heieis et al., 2023). The complete list of antibodies is supplied in Table 5. Cells were analysed on an ID7000^TM^ 7-laser Spectral Cell Analyzer (Sony Cooperation).

### Liver glycogen assay

To access the glycogen content of perinatal liver tissue the Liver Glycogen Assay Kit from Abcam (ab65620) was used. The kit components were stored and dissolved following the manufacturerś instructions. The used liver samples were snap-frozen in liquid nitrogen and stored at −80 °C until the experiment. 10 mg of liver tissue per sample were weighed in and the serum samples were diluted (1:25-1:100). The liver tissue was then washed shortly with cold PBS and put into 500 μL of ddH2O. Homogenization was performed with a pestle on ice and the homogenates were boiled at 95 °C for 10 min to inactivate enzymes. Further, the boiled homogenates were centrifuged at 4 °C, maximal speed, and the supernatant was collected and stored at −20 °C until the assay was performed. The main assay procedure was performed as described in the manufactures’ instructions. All samples and the glycogen standard curve were measured in technical duplicates. The plate was measure immediately after the last step with a Tecan Infinite M200 microplate reader at Ex/Em = 535/587 nm. For analysis, the measured fluorescence from each sample was corrected with a glucose control (no addition of glucoamylase) to account for intrinsic differences in hepatic/systemic glucose levels.

### Igf1 ELISA

To determine the amount of Igf1 present in the perinatal liver and serum the Quantitative ELISA Mouse/Rat Igf1 Kit Liver Glycogen Assay Kit from R&D Systems (MG100) was used. The kit components were stored and dissolved following the manufacturerś instructions. The samples were snap-frozen on liquid nitrogen and stored at −80 °C until the experiment was performed. 10 mg liver tissue was homogenized in 500 μL ddH2O on ice and cell membranes broken through two repeated freeze-thawing cycles. The liver homogenates were centrifuged at 5,000g (5 min, 4 °C) and the supernatant was collected. The serum samples were diluted 1:100 before analysis. For the ELISA, the manufacturer’s instructions were followed. The optical density was determined using a Tecan Infinite M200 plate reader at 450 nm with correction at 540 nm.

### Lipidomics analysis

To evaluate differences in hepatic lipid metabolism tandem mass spectrometry of extracted lipids was performed. For this purpose, 10 mg liver tissue was homogenized in 500 μL ddH2O on ice. Then, 50 μL of the homogenate was transferred into a fresh Eppendorf tube and 500 μL Extraction Mix (CHCl_3_/MeOH 1/5 containing the following internal standards: 210 pmol PE(31:1), 396 pmol PC(31:1), 98 pmol PS(31:1), 84 pmol PI(34:0), 56 pmol PA(31:1), 51 pmol PG (28:0), 28 pmol CL(56:0), 39 pmol LPA (17:0), 35 pmol LPC(17:1), 38 pmol LPE (17:0), 32 pmol Cer(17:0), 99 pmol, SM(17:0), 55 pmol GlcCer(12:0), 14 pmol GM3 (18:0-D3), 339 pmol TG(50:1-d4), 111 pmol, CE(17:1), 64 pmol DG(31:1), 103 pmol MG(17:1), 724 pmol Chol(d6) and 45 pmol Car(15:0) if everything is correct here) was added. After 2 min of sonication in a bath sonicator, the samples were spun at 20,000 g for 2 min. The supernatant was collected in a new Eppendorf tube and 200 μL chloroform and 750 μL of 1 M NH_4_Ac in ddH_2_O were added. Following quick manual shaking, the samples were centrifuged at 20,000 g for 2 min again. The upper phase was carefully removed, and the lower phase was transferred into a new Eppendorf tube. The solvent was evaporated using a SpeedVac Vacuum Concentrator at 45 °C for 20 min. The dried lipids were dissolved in 500 μL Spray Buffer (Isopropanol, Methanol, ddH2O (all MS grade), 10 mM ammonium acetate, 0.1 % acetic acid by sonication for 5 min. Until measurement with a Thermo Q Exactive ™ Plus (Thermo Scientific) using positive mode, the samples were stored at −20 °C. Before the acquisition, the samples were sonicated for 5 min. For downstream analysis, the raw spectral data was converted to .mzml files and loaded into the LipidXplorer software (Herzog et al., 2012). Using both the sample lipid and the previously added internal standard it calibrates the mass spectra and also discriminates based on the mass of the different lipid species. The sample lipid concentration in pmol was calculated referring to the intensity of the internal standard peak for each sample and its known concentration in the added internal standard. Samples with a high deviation of the internal standard were excluded from analysis. The overall abundance of the respective lipid class was obtained by summarizing the species’ amounts. Lipidomics datasets are accessible via Metabolomics Workbench (Sud et al., 2016) under the study IDs ST003614 (*WT^Spi1^/KO^Spi1^*) and ST003702 (*WT^Csf1r^/KO^Csf1r^*).

### Phosphoproteome analysis

Liver tissues were lysed in 4% sodium deoxycholate (SDC) and 100 mM Tris-HCl (pH 8.5), followed by immediate boiling at 95 °C for 5 minutes. The lysates were then sonicated for 20 minutes at 4 °C. Protein concentrations were determined using the BCA protein assay, and samples were adjusted to 1 mg of total protein. Samples were reduced and alkylated using 10 mM tris(2-carboxyethyl)phosphine (TCEP) and 40 mM 2-chloroacetamide (CAM). Digestion was performed overnight at 37 °C using lysC and trypsin at an enzyme-to-protein ratio of 1:100 (wt/wt). The digested peptides were treated with isopropanol (final concentration 50%), trifluoroacetic acid (TFA, final concentration 6%), and monopotassium phosphate (KH₂PO₄, final concentration 1 mM). Samples were mixed at 1500 rpm for 30 seconds, cleared by centrifugation, and the supernatants were incubated with TiO₂ beads (Titansphere Phos-TiO Bulk, GL Sciences) at a bead-to-protein ratio of 12:1 (wt/wt) for 5 minutes at 40 °C. The beads were washed five times with 60% isopropanol and 5% TFA, and phosphopeptides were eluted using 40% acetonitrile (ACN) and 15% ammonium hydroxide (NH₄OH, 25%, HPLC grade). The eluates were collected by centrifugation into clean PCR tubes and concentrated using a SpeedVac for 20 minutes at 45 °C. Finally, phosphopeptides were desalted on SDB-RPS StageTips and resuspended in 7 μl of 2% ACN and 0.3% TFA for LC-MS/MS analysis. Phosphopeptide samples were analyzed using an EASY-nLC 1000 HPLC system (Thermo Fisher Scientific) coupled to a Q Exactive HFX mass spectrometer (Thermo Fisher Scientific) via a nano-electrospray ion source. The samples were loaded onto a 50-cm column with a 75 μm inner diameter, packed in-house with C18 1.9 μM ReproSil particles (Dr. Maisch GmbH). The column temperature was maintained at 50 °C using a custom-built column oven. Phosphopeptides were separated using a buffer system consisting of buffer A (0.1% formic acid) and buffer B (0.1% formic acid, 80% ACN). A 120-minute gradient was used for elution, starting at ?% buffer B and increasing stepwise to ?% inminutes, ?% inminutes, and ?% inminutes at a flow rate of 300 nl/min. Phosphopeptides were analyzed using a data-independent acquisition (DIA) MS method. This included one full scan (300–1650 m/z, R = 60,000 at 200 m/z) with a target of 3 × 10⁶ ions, followed by 32 windows with a resolution of 30,000 where precursor ions were fragmented with higher-energy collisional dissociation (stepped collision energy 25%, 27.5%, and 30%) and analyzed with an AGC target of 3e^6^ ions and a maximum injection time at 54 ms in profile mode using positive polarity.

Raw MS data files were processed using Spectronaut software (version 14.3.200701.47784). Mass spectra were searched against the mouse UniProt FASTA database (July 2019, 63,439 entries) with a false discovery rate (FDR) of less than 1% at the protein and peptide levels. Database search parameters allowed a minimum peptide length of 7 amino acids and a maximum of two missed cleavages. Variable modifications included phosphorylation on serine, threonine, and tyrosine residues, N-terminal acetylation, and methionine oxidation, while cysteine carbamidomethylation was set as a fixed modification. All bioinformatics analyses were conducted in R (v4.0.4). Quantified phosphosites were filtered to include only those with at least three valid values across biological replicates in at least one condition. Missing values were imputed using a Gaussian distribution with a width of 0.3 and a downshift of 1.8 standard deviations of the measured values. Differentially regulated phosphosites were identified using a Student’s t-test with a permutation-based FDR of 5%. A 1D annotation enrichment analysis was applied to identify systematic enrichment or depletion of annotations and pathways (FDR < 5%).

### PAS staining

Periodic acid-Schiff’s with hematoxylin (PAS-H) staining was performed on paraffin-embedded tissue sections (5 µm thick) to analyze glycogen storage in neonatal liver tissues. Tissue sections were incubated at 65 °C for 30 minutes to melt the paraffin. Rehydration was performed by passing the sections through a series of steps, each for 3 minutes: 2 changes of xylene, followed by 100%, 95%, 90%, 80%, and 70% ethanol, and finally distilled water. Rehydrated sections were incubated in periodic acid solution (Carl Roth, cat. No: HP00) for 10 minutes to oxidize glycogen, rinsed in tap water for 3 minutes, and briefly rinsed in distilled water. The sections were placed in Schiff’s reagent (Carl Roth, cat. No: X900.2) for 15 minutes. Following staining, sections were washed in running tap water for 10 minutes then briefly rinsed in distilled water. Counterstaining was performed using hematoxylin (Carl Roth, cat. No: T865) for 45 seconds to visualize nuclei. Excess hematoxylin was removed by washing in tap water for 3 minutes, followed by a brief rinse in distilled water. Sections were passed through distilled water, 70%, 80%, 90%, 95%, and 100% ethanol, each for 3 minutes, and then cleared in 2 changes of xylene. Sections were mounted with Entellan (Merck, cat. No: 107961) and covered with glass coverslips. Stained sections were examined and recorded with a light microscope. For analysis of acquired images Fiji analysis software (v2.1.0/1.53c) was used. First, the color threshold was set for the images and they were converted to 8-bit images. After setting an auto threshold, they were reverted, and the color deconvoluted. Signal intensity for the PAS image plane was measured, and all values were normalized using the respective WT reference.

### Transmission electron microscopy

Transmission electron microscopy of mouse liver samples was performed as described previously for other organs (Fazio et al., 2022; Welz et al., 2022). To quantify cristae area of mitochondria, images were acquired at a nominal magnification of 20k and mitochondria were manually singled out using ImageJ. Subsequently, we applied an in house developed script utilizing ImageMagick (www.imagemagick.org). This script binarized the mitochondrial images applying the same threshold value for all samples. We then counted the black pixels in each mitochondrion as a measure for mitochondrial mass. To obtain the cristae area, we subtracted the black pixels from the overall mitochondrial pixel number. To calculate the cristae area per mitochondria we divided the cristae area through the overall area.

### Perinatal liver culture and metabolic tracing

Newborn pups were sacrificed immediately after birth by decapitation, and the liver was collected in ice-cold culture buffer (Williams Medium supplemented with 10% FCS, 1% L-glutamine, and 1% Penicillin-Streptomycin). For each replicate, 2–3 mg of the largest liver lobe was carefully dissected and placed into a 48-well plate containing ice-cold culture buffer. All subsequent steps were carried out in a sterile hood to maintain sterility. Each liver piece was transferred onto a porous cell culture insert (0.4 µm pore size), which was placed in a well of a 24-well plate containing 400 µL of prewarmed culture buffer. The plate was incubated at 37 °C with 5% CO₂ for 2 hours. This incubation period allowed for the simultaneous genotyping of the pups to ensure that only WTSpi1 and KOSpi1 samples were included in the tracing experiment, excluding HETSpi1 samples. After genotyping, the inserts with the liver tissue pieces were transferred to a new 24-well plate containing 400 µL of prewarmed tracer medium (Williams Medium without glucose, supplemented with 10% FCS, 1% L-glutamine, and 25 mM C¹³-labeled glucose). The samples were incubated for 4 hours at 37 °C with 5% CO₂. Following the incubation, 100 µL of medium was collected and snap-frozen. The liver tissue pieces were briefly washed with 0.9% NaCl and then placed in 250 µL of methanol pre-cooled to −20 °C. To each sample, 250 µL of pre-cooled MS-grade water containing 1 µg/mL D6-glutaric acid (used as an internal standard) was added, and the tissue was homogenized. The homogenate was transferred to a fresh tube and stored at −80 °C until further processing. Metabolites were derivatized using a Gerstel MPS with 15 μl of 2% (w/v) methoxyamine hydrochloride (Thermo Scientific) in pyridine and 15 μl N-tertbutyldimethylsilyl-N-methyltrifluoroacetamide (MTBSTFA) with 1% tert-butyldimethylchlorosilane (Regis Technologies). Derivatives were measured by GC/MS with a 30 m DB-35MS + 5 m Duraguard capillary column (0.25 mm inner diameter, 0.25 µm film thickness) equiped in an Agilent 7890B gas chromatograph (GC) connected to an Agilent 5977A mass spectrometer (MS). The GC oven temperature was held at 80 °C for 6 min and subsequently increased at 6 °C per min until reaching 280 °C where it was held for 10 min. The quadropole was held at 150 °C. The MS source operated under electron impact ionization mode at 70 eV and was held at 230°C.

Full scan (70–800 *m*/*z*, 3.9 scans per second) as well as targeted ion chromatogram measurements were conducted for pyruvate (174, 175, 176, 177, 178, 179; 10 scans per second), lactate (261, 262, 263, 264, 265, 266, 267; 10 scans per second), alanine (260, 261, 262, 263, 264, 265; 10 scans per second), citrate (591, 592, 593, 594, 595, 596, 597, 598, 599, 600; 10 scans per second), malate (419, 420, 421, 422, 423, 424, 425, 426; 10 scans per second) and aspartate (418, 419, 420, 421; 10 scans per second). All chromatograms were analyzed with MetaboliteDetector (Hiller et al., 2009). ^13^C-glucose tracing datasets are accessible via Metabolomics Workbench (Sud et al., 2016) under the project number ST003615.

### Bulk-RNA sequencing

Cells were stored into Trizol and stored at −80°C. The cDNA library for sequencing was prepared following the SMART-Seq2 protocol. mRNA was isolated, primed using poly-T oligonucleotides, and then converted into cDNA via SMART reverse transcription. Pre-amplification was performed using SMART ISPCR, followed by fragmentation with the Nextera XT DNA Library Preparation kit (Illumina), amplification, and indexing. Library fragments were then selected by size (300-400 bp) and purified using SPRIBeads (Beckman-Coulter). Size distribution of the cDNA libraries was analyzed using the Agilent high-sensitivity D5000 assay on the Tapestation 4200 system (Agilent). Quantification of cDNA libraries was completed with a Qubit high-sensitivity dsDNA assay (Thermo Fisher). Sequencing employed a 75 bp single-end configuration on the NextSeq500 system (Illumina), using the NextSeq 500/550 High Output Kit v2.5. Kallisto pseudo-alignment was applied to quantify transcript abundances from bulk RNA-seq data (Bray et al., 2016). The files were processed using Kallisto for transcript quantification, with the Gencode M16 mouse annotation (https://www.gencodegenes.org/mouse/release_M16.html) applied to adjust for library size based on the average transcript length. Read counts were normalized with DESeq2’s default size factor estimation, which corrects for sequencing depth across samples. Low-expressed genes, defined as those with fewer than 10 total counts, were removed to enhance signal detection. Statistical significance in gene expression was determined using the Wald test for pairwise comparisons (default method in DESeq2) Adjusted p-values were calculated using the Benjamini-Hochberg method to control the false discovery rate (FDR). Genes with an adjusted p-value < 0.05 and |log2fold change| > 1.3 were considered significantly differentially expressed. Normalized count values for individual genes were visualized to assess expression differences. Genes were ranked by differential expression for each condition, and the resulting ranked gene list was used for gene set enrichment analysis (GSEA) on selected gene sets.

### Single-cell RNA-sequencing library preparation and analysis

Single-cell suspensions were prepared as previously described. The cells were then spun for 3 minutes at 50 g to enrich hepatocytes. Both hepatocytes and liver immune cells were utilized for single-cell analysis. To ensure that hepatocytes are appropriately sized to fit into the wells of the Seq-Well array, the cells were fixed to induce controlled shrinkage. Hepatocytes were resuspended in PBS and fixed by the slow, dropwise addition of ice-cold methanol at a 1:5 ratio (PBS:methanol) while gently stirring the solution between additions. The suspension was incubated at −20°C for 30 min, followed by 5 min on ice. Subsequently, the fixed cells were centrifuged at 1000 g for 5 min at 4°C, and the supernatant was carefully discarded. The hepatocytes were resuspended in 500 µl of rehydration buffer containing 3× SSC (Sigma, Cat. S6639-1L), 1 mM Dithiothreitol (Thermo Fisher Scientific, Cat. R0862), 0.2 U/µl RNase inhibitor (Thermo Fisher Scientific, Cat. AM2696), and 2 mM flavopiridol (Sigma, Cat. F3055). Cell loading, barcoding and library preparation primarily followed the Seq-Well S3 protocol (Hughes et al., 2020), with two arrays per sample. Seq-Well arrays were set up as described by (Gierahn et al., 2017). Each array was loaded with approximately 110,000 barcoded mRNA capture beads (ChemGenes, Cat: MACOSKO-2011-10) and 30,000 cells. Following cell loading, cells were lysed, mRNA captured, and cDNA synthesis was performed. For whole transcriptome amplification, beads from each array were divided into 18-24 PCR reactions with approximately 3,000 beads per reaction (95°C for 3 min, 4 cycles of 98°C for 20 s, 65°C for 45 s, 63°C for 30 s, 72°C for 1 min; followed by 16 cycles of 98°C for 20 s, 67°C for 45 s, 72°C for 3 min; final extension at 72°C for 5 min) using the KAPA HiFi Hotstart Readymix PCR Kit (Kapa Biosystems, Cat: KK2602) and SMART PCR Primer (AAGCAGTGGTATCAACGCAGAGT). Pooled PCR reactions (6-8 per pool) were purified using AMPure XP SPRI Reagent (Beckman Coulter) with sequential 0.6x and 1x volumetric ratios. For library tagmentation and indexing, 200 pg of DNA from each purified WTA pool was tagmented with the Nextera XT DNA Library Preparation Kit (8 min at 55°C), followed by Tn5 transposase neutralization (5 min at RT). Illumina indices were then attached to the tagmented products (72°C for 3 min, 98°C for 30 s; 16 cycles of 95°C for 10 s, 55°C for 30 s, 72°C for 1 min; final extension at 72°C for 5 min). The library products were purified using AMPure XP SPRI Reagent at 0.6x and 1x volumetric ratios. The final library quality was assessed using a High Sensitivity D5000 assay on a Tapestation 4200 (Agilent) and quantified with the Qubit high-sensitivity dsDNA assay (Invitrogen). Seq-Well libraries were pooled equimolarly and clustered at a 1.25 nM concentration with 10% PhiX on a NovaSeq6000 system (S2 flow cell, 100 bp v1.5 chemistry). Sequencing was paired-end, using a custom Drop-Seq Read 1 primer for 21 cycles, 8 cycles for the i7 index, and 61 cycles for Read 2. Single-cell data were demultiplexed using bcl2fastq2 (v2.20). Fastq files from Seq-Well were processed in a snakemake-based pre-processing pipeline (v0.31, available at https://github.com/Hoohm/dropSeqPipe) that utilizes Drop-seq tools provided by the McCarroll lab (Macosko, Basu et al. 2015). STAR alignment within the pipeline was performed using the murine GENCODE reference genome and transcriptome (mm10 release vM16; Team 2014). To analyze liver cells, UMI-corrected expression matrices were processed in R using Seurat (v.4.1.0)(Hafemeister and Satija, 2019) From an initial dataset of 407,803 barcodes and 43,126 genes, only protein-coding genes were retained, reducing the dataset to 16,131 genes. Quality control measures (Luecken and Theis, 2019) identified 27,258 high-quality cells. Ambient RNA contamination was corrected using SoupX (Young and Behjati, 2020) further refining the dataset to 27,235 cells. Data normalization and scaling were performed with SCTransform (Hafemeister and Satija, 2019) dimensionality reduction was achieved through Principal Component Analysis (PCA). The first 21 principal components were selected for downstream analyses, including UMAP visualization and the construction of a Shared Nearest Neighbor (SNN) graph. Clustering was performed using the Louvain algorithm (Traag et al., 2019), with resolution determined by Clustree v.0.4.3 (Zappia and Oshlack, 2018) and NbClust v.3.0 (Charrad et al., 2014) analyses. Marker genes for each cluster were identified using a Wilcoxon rank sum test (Haynes, 2013), applying a log_2_ fold-change threshold of 0.25 and requiring expression in at least 10% of cells. Cluster annotation was based on canonical marker genes. To investigate functional phenotypes in hepatocytes, a Wilcoxon rank sum test was performed without thresholds to identify all DEGs between *WT^Spi1^* and *KO^Spi^* genes were ranked by average log_2_ fold-change for each genotype and the top 100 genes were used for GOEA (Yu et al., 2012; Haynes, 2013) and the Mouse annotation database (org.Mm.eg.db) v.3.12.0. GSEA was conducted using ClusterProfiler to identify enriched biological processes from the Gene Ontology (GO) database (Ashburner et al., 2000). Statistical significance of enriched terms was evaluated using a hypergeometric test (Yu et al., 2015) with Bonferroni correction for multiple comparisons (Yu et al., 2015, 2012; Haynes, 2013). Terms with adjusted p-values below 0.05 were considered significantly enriched. Developmental trajectories of hepatocyte subclusters were inferred using Monocle3 v1.0.0 (Cao et al., 2019). For trajectory construction, hepatocytes were subsetted from the liver scRNAseq dataset and the first 30 dimensions of the scRNA-seq data were utilized to generate a UMAP focused exclusively on hepatocytes. A connection matrix was computed using a modified partitioned approximate graph abstraction (PAGA) algorithm (Wolf et al., 2017), followed by significance testing of connections between clusters identified through PAGA and Louvain clustering. Subsequently, a principal graph was constructed in the low-dimensional space using a modified SimplePPT algorithm (Mao et al., 2015). Pseudotime progression was determined by designating the node within most immature hepatocyte cluster 0 as the root node based on hepatocyte maturation marker expression, enabling the inference of developmental relationships among hepatocyte subclusters. Intercellular communication between hepatocytes and Kupffer cells was analyzed following the methods described by Jin et al. (Jin et al., 2021), utilizing CellChat v1.1.3. The Seurat object was split into three datasets based on genotypes. Each dataset was converted into a CellChat object using normalized and log1p-transformed expression matrices (Jin et al., 2021). The dataset was then subset to include only genes associated with known signaling pathways. Overexpressed ligand and receptor genes were identified using the Wilcoxon rank sum test, without applying a p-value threshold. Noise was mitigated by calculating the average expression of each gene for each cell group, with weighted summation of gene expression quantiles. The prediction of gene-gene interactions relied on Protein-Protein Interaction (PPI) networks sourced from STRINGDb (Szklarczyk et al., 2019) under the assumption that physical interactions between ligands and receptors follow the law of mass action. To achieve this, signaling gene expression profiles were projected onto the PPI network using random walk network propagation (Cowen et al., 2017). The interaction probability (or strength) was modeled using the projected data, applying a trimming fraction of 0.01. Communication pathways between cell groups were identified through permutation testing. Additionally, social network analysis (SNA) tools from the sna package (Butts, 2008) were used to calculate information flow metrics, including out-degree, in-degree, flow betweenness, and information centrality, to evaluate intercellular communication.

## Code and Data Availability

All steps, including cleaning, dimensionality reduction, clustering, differential gene expression (DEG) testing, and Gene Ontology Enrichment Analysis (GOEA), were conducted using the docker image alefrol94/scrnaseq.analysis:reticulate. Additional packages were managed and tracked using the renv package (v0.14.0). Trajectory analysis of hepatocyte subsets, performed with Monocle3 (Cao et al., 2019), utilized the docker image jsschrepping/r_docker:jss_R403_S4cran. The complete analysis code can be made available upon request. scRNA-seq datasets are accessible via the Gene Expression Omnibus (GEO) repository under the accession number GSE285047. The baulk RNA-seq raw transcriptome files and count data have been deposited in NCBI’s Gene Expression Omnibus (GEO) and are accessible under accession number GSE237408.

## Experimental design, quantification and statistical analysis

For the design of experiments, scientists were blinded to the experimental groups whenever possible, as the animals exhibited no overt phenotype at the time of sample processing. Blinding was applied during various stages, such as histological and transmission electron microscopy (TEM) analyses, where subjective interpretation could introduce bias. However, full blinding was not feasible in all assays. For example, genotyping PCR and certain analyses, such as flow cytometry, revealed clear phenotypic differences, which could not be masked from experimenters. To account for variability between experimental runs, we normalized data to the mean value(s) of littermate controls collected on the same day. This normalization was particularly important for measurements such as the mean fluorescence intensity (MFI) of flow cytometry markers and most metabolic analyses, ensuring consistency and comparability across experiments. Every reported n value represents the number of biologically independent replicates. No statistical methods were used to predetermine sample sizes, but sample numbers were based on standards in the field and experimental feasibility. All statistical analyses, except for those related to sequencing data, were conducted using GraphPad Prism software (versions 5–8; GraphPad Software, RRID:SCR_002798). For comparisons between two groups, Mann-Whitney U-tests, paired/unpaired Student’s t-tests (e.g., paired tests for histological analysis due to variability in staining intensity across experiments), were performed depending on data distribution and experimental design. Wald test was performed on the fold-change gene expression of KCs and KC-like cells for pairwise comparisons. Differences were considered statistically significant at p values less than 0.05, with ns denoting non-significance (p > 0.05). Each dataset is representative of at least two independent experiments, with a minimum of three animals per group. Detailed descriptive statistics, the specific statistical tests performed, and the number of samples analyzed are provided in the corresponding figure legends.

## Author contributions

NM: Formal analysis, Investigation, Methodology, Validation, Visualization, Writing – original draft, Writing – review & editing. DH: Formal analysis, Investigation, Methodology, Software, Validation, Visualization, Writing – original draft, Writing – review & editing. AF: Data curation, Software, Visualization, Writing – review & editing. IS: Formal analysis, Investigation, Writing – review & editing. FN: Data curation, Formal analysis, Visualization, Writing – review & editing. JC: Data curation, Formal analysis, Visualization, Writing – review & editing. NB-S: Investigation, Writing – review & editing. MFV: Investigation, Writing – review & editing. MY: Investigation, Writing – review & editing. PA: Data curation, Formal analysis, Visualization, Writing – review & editing. LB: Formal analysis, Writing – review & editing; MB: Data curation, Writing – review & editing. CT: Funding acquisition, Resources, Supervision, Writing – review & editing. FM: Funding acquisition, Resources, Supervision, Writing – review & editing. KH: Funding acquisition, Resources, Supervision, Writing – review & editing. MDB: Funding acquisition, Resources, Supervision, Writing – review & editing. EM: Conceptualization, Formal analysis, Funding acquisition, Methodology, Resources, Supervision, Visualization, Writing – original draft, Writing – review & editing.

## Acknowledgements

We thank Cornelia Cygon for the support in the lab. We thank Andrea Eichhorn and Elke Kretzschmar for their help with the TEM images. We thank Yasuhiro Kobayashi for providing *Tnfrsf11a^Cre^* mice and Florent Ginhoux for providing the *Ms4a3^FlpO^* mice. The work in the labs was supported by the following grants: Funded by the Deutsche Forschungsgemeinschaft (DFG, German Research Foundation) under Germany’s Excellence Strategy-EXC2151-390873048 (to EM, MDB, FM, CT, LB), Project-ID 432325352 – SFB 1454 (to EM, MDB, KH, FM, CT), GRK1873/2 (to NM, EM), IRTG program GRK2168 (Project number 272482170 to MDB), FOR5547 – Project-ID 503306912 (to EM), ImmuDiet (Project ID 432325352 to LB). European Research Council (ERC) under the European Union’s Horizon 2020 research and innovation program (Grant Agreement No. 851257 to EM, Grant Agreement No. 101163024, POLIS, to LB). We would like to thank the Flow Cytometry Core Facility of the Mathematical and Natural Sciences Faculty and Flow the Cytometry Core Facility Medical Faculty at the University of Bonn for providing support and instrumentation funded by the Deutsche Forschungsgemeinschaft (DFG, German Research Foundation) - Project numbers 341039622, 144734146, 471514137, 01EO2107 (BMBF).

---

**Figure S1.**
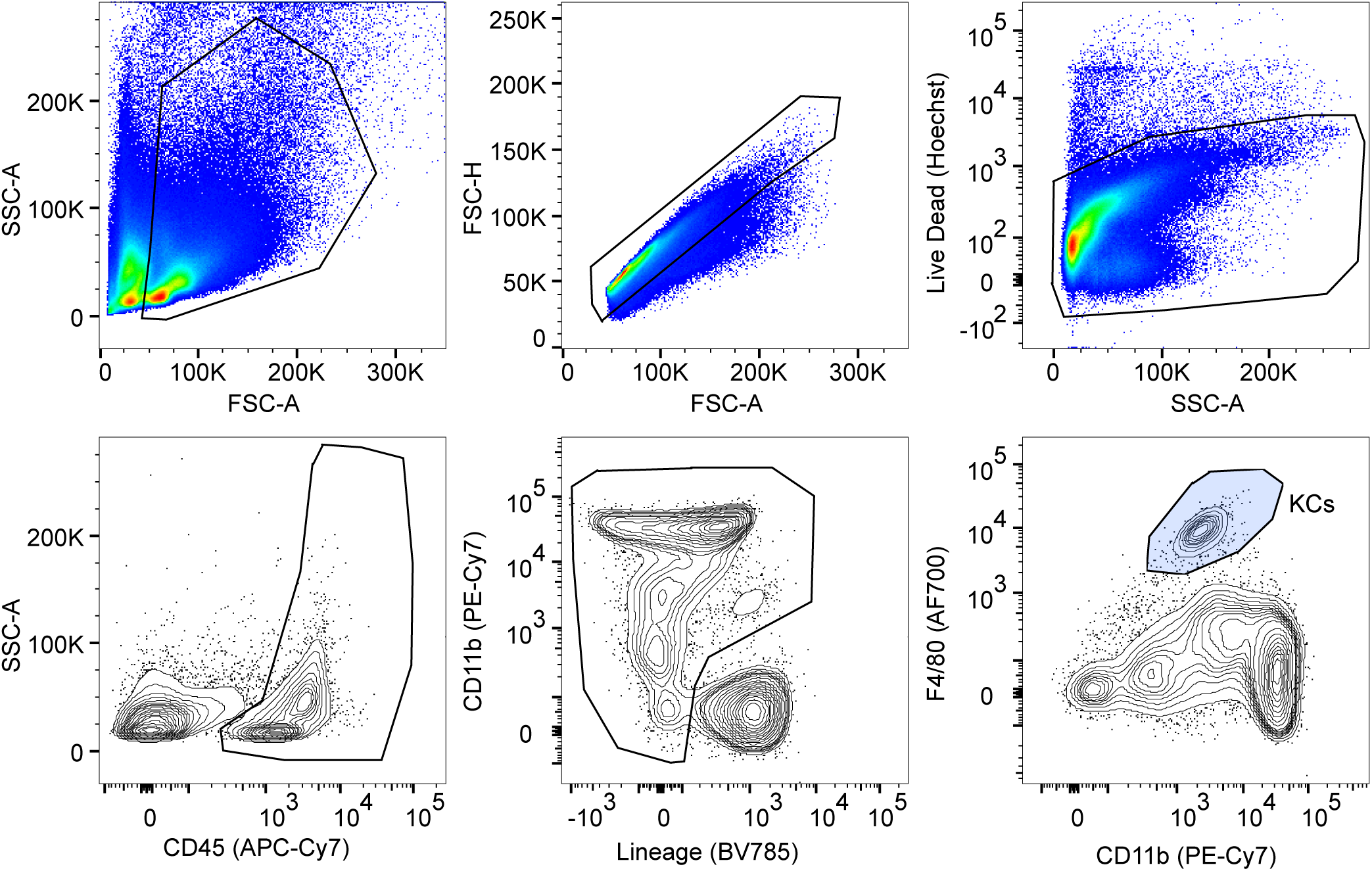
Flow cytometry gating for Kupffer Cells. Gating strategy to identify macrophage clusters in E12.5, E14.5, and P0 livers.

**Figure S2.**
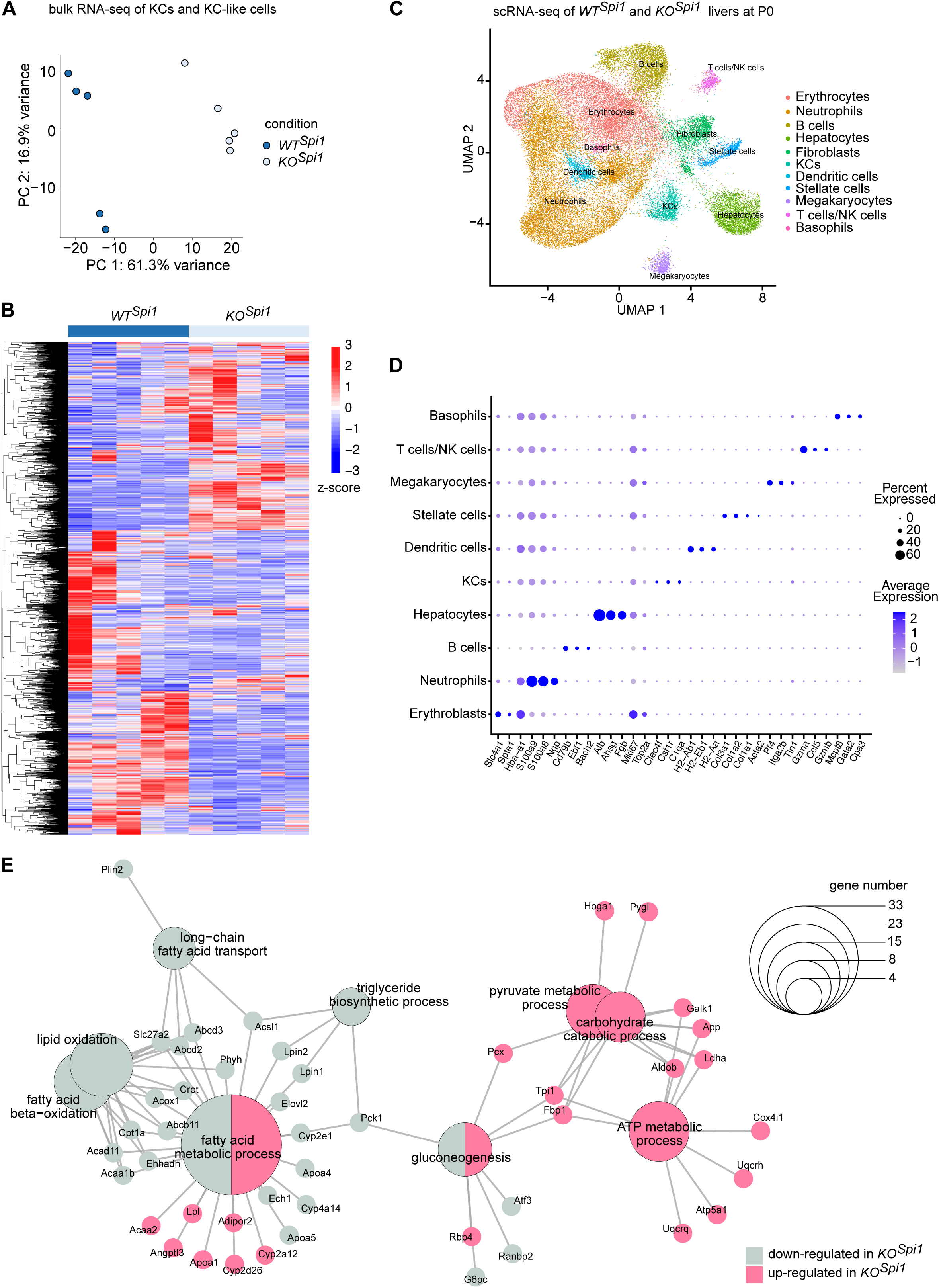
Bulk RNA-Seq and scRNA-Seq analysis of the *KO^Spi1^* mouse model. **(A)** PCA visualization of the *WT^Spi^* and *KO^Spi^*KC and KC-like macrophages, respectively. **(B)** Heatmap for the most variable genes between *WT^Spi^* and *KO^Spi^* macrophages. **(C)** UMAP visualization of the annotated single cell clusters of *WT^Spi^* and *KO^Spi^* livers. **(D)** Dotplot showing canonical marker genes to identify specific clusters. **(E)** Network visualization of gene ontology enrichment analysis of DEGs on hepatocyte cluster 2 showing the pathways and the involved genes.

**Figure S3.**
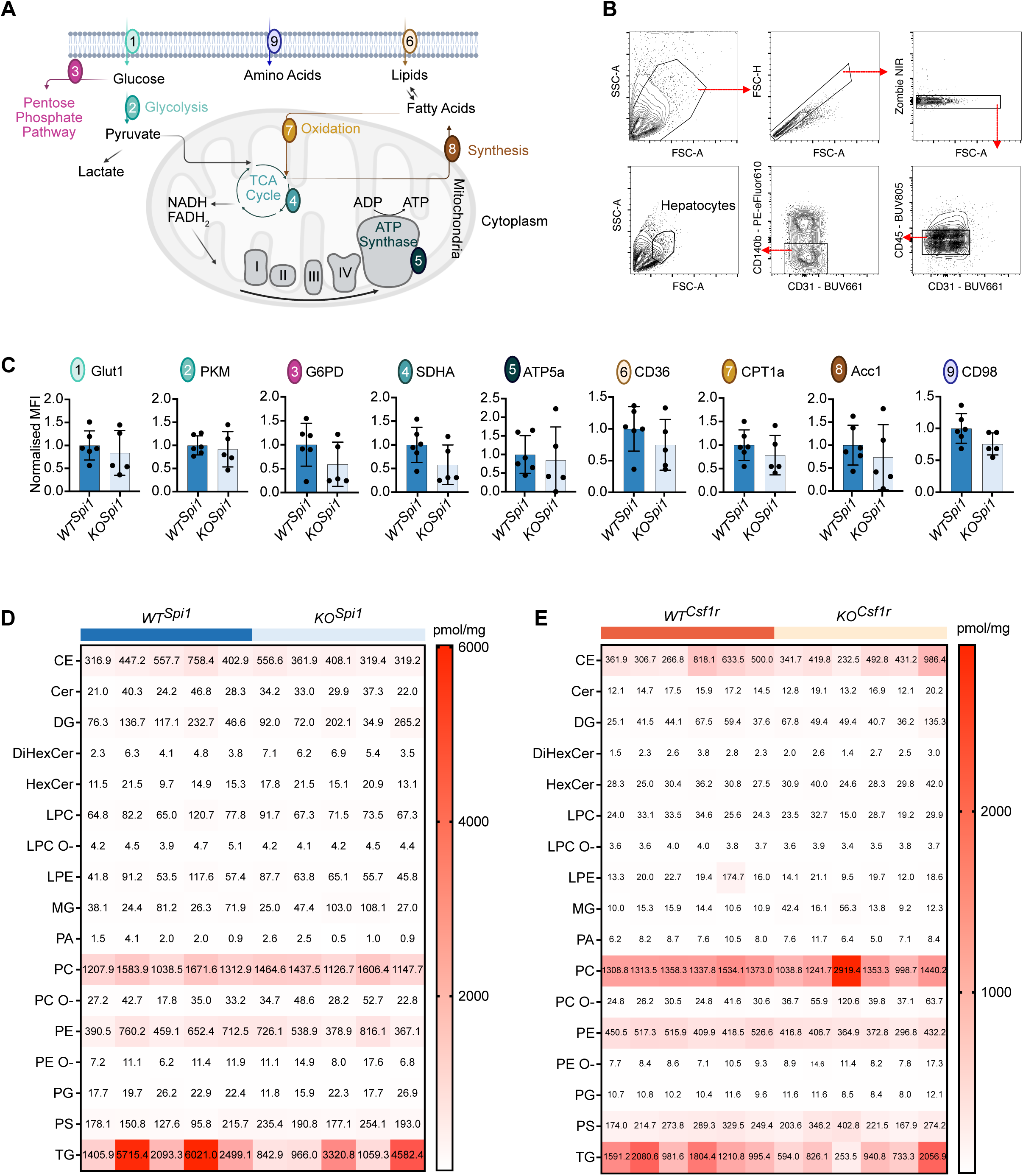
Metabolic phenotyping of the *KO^Spi1^* mouse model: **(A)** Schematic of metabolic targets for flow cytometry-based analysis. Created with BioRender.com. **(B)** Representative gating strategy of hepatocytes. **(C)** Normalized expression of metabolic targets in hepatocytes at P0 from *WT^Spi^* and *KO^Spi^*. n = 5-6 per genotype from 2 litters. Barplot presented as mean ± SD. Mann-Whitney test. **(D-E)** Lipid species abundance in *WT^Spi^* and *KO^Spi^* (D) and *WT^Csf1r^* and *KO^Csf1r^* livers (E). n = 5-6 per genotype from more than 3 litters per mouse model. CE: Cholesteryl Ester, Cer: Ceramide, DG: Diacylglycerol, DiHexCer: Di-Hexosylceramide, HexCer: Hexosylceramide, LPC: Lysophosphatidylcholine, LPC O-: Alkyl-Lysophosphatidylcholine, LPE: Lysophosphatidylethanolamine, MG: Monoacylglycerol, PA: Phosphatidic Acid, PC: Phosphatidylcholine, PC O-: Alkyl-ether phosphatidylcholine, PE: Phosphatidylethanolamine, PE O-: Alkyl-Phosphatidylethanolamine, PG: Phosphatidylglycerol, PS: Phosphatidylserine, TG: Triacylglycerol.

## References

Araujo David, B., J. Atif, F. Vargas E Silva Castanheira, T. Yasmin, A. Guillot, Y. Ait Ahmed, M. Peiseler, J.W. Hommes, L. Salm, M.A. Brundler, B.G.J. Surewaard, W. Elhenawy, S. MacParland, F. Ginhoux, K. McCoy, and P. Kubes. 2024. Kupffer cell reverse migration into the liver sinusoids mitigates neonatal sepsis and meningitis. Sci Immunol. 9:eadq9704. doi:10.1126/SCIIMMUNOL.ADQ9704.

Ashburner, M., C.A. Ball, J.A. Blake, D. Botstein, H. Butler, J.M. Cherry, A.P. Davis, K. Dolinski, S.S. Dwight, J.T. Eppig, M.A. Harris, D.P. Hill, L. Issel-Tarver, A. Kasarskis, S. Lewis, J.C. Matese, J.E. Richardson, M. Ringwald, G.M. Rubin, and G. Sherlock. 2000. Gene Ontology: tool for the unification of biology. Nature Genetics 2000 25:1. 25:25–29. doi:10.1038/75556.

Bray, N.L., H. Pimentel, P. Melsted, and L. Pachter. 2016. Near-optimal probabilistic RNA-seq quantification. Nature Biotechnology 2016 34:5. 34:525–527. doi:10.1038/nbt.3519.

Cao, J., M. Spielmann, X. Qiu, X. Huang, D.M. Ibrahim, A.J. Hill, F. Zhang, S. Mundlos, L. Christiansen, F.J. Steemers, C. Trapnell, and J. Shendure. 2019. The single-cell transcriptional landscape of mammalian organogenesis. Nature 2019 566:7745. 566:496–502. doi:10.1038/s41586-019-0969-x.

Charrad, M., N. Ghazzali, V. Boiteau, and A. Niknafs. 2014. NbClust: An R Package for Determining the Relevant Number of Clusters in a Data Set. J Stat Softw. 61:1–36. doi:10.18637/JSS.V061.I06.

Cowen, L., T. Ideker, B.J. Raphael, and R. Sharan. 2017. Network propagation: a universal amplifier of genetic associations. Nature Reviews Genetics 2017 18:9. 18:551–562. doi:10.1038/nrg.2017.38.

Cox, N., L. Crozet, I.R. Holtman, P.-L. Loyher, T. Lazarov, J.B. White, E. Mass, E.R. Stanley, O. Elemento, C.K. Glass, and F. Geissmann. 2021. Diet-regulated production of PDGFcc by macrophages controls energy storage. Science (1979). 373:eabe9383. doi:10.1126/science.abe9383.

Diehl, K.L., J. Vorac, K. Hofmann, P. Meiser, I. Unterweger, L. Kuerschner, H. Weighardt, I. Förster, and C. Thiele. 2020. Kupffer Cells Sense Free Fatty Acids and Regulate Hepatic Lipid Metabolism in High-Fat Diet and Inflammation. Cells. 9. doi:10.3390/CELLS9102258.

Elmore, M.R.P., L.A. Hohsfield, E.A. Kramár, L. Soreq, R.J. Lee, S.T. Pham, A.R. Najafi, E.E. Spangenberg, M.A. Wood, B.L. West, and K.N. Green. 2018. Replacement of microglia in the aged brain reverses cognitive, synaptic, and neuronal deficits in mice. Aging Cell. 17:e12832. doi:10.1111/ACEL.12832.

Fazio, A., D. Bordoni, J.W.P. Kuiper, S. Weber-Stiehl, S.T. Stengel, P. Arnold, D. Ellinghaus, G. Ito, F. Tran, B. Messner, A. Henning, J.P. Bernardes, R. Häsler, A. Luzius, S. Imm, F. Hinrichsen, A. Franke, S. Huber, S. Nikolaus, K. Aden, S. Schreiber, F. Sommer, G. Natoli, N. Mishra, and P. Rosenstiel. 2022. DNA methyltransferase 3A controls intestinal epithelial barrier function and regeneration in the colon. Nat Commun. 13. doi:10.1038/S41467-022-33844-2.

Fellinger, P., H. Beiglböck, G. Semmler, L. Pfleger, S. Smajis, C. Baumgartner, M. Gajdosik, R. Marculescu, G. Vila, Y. Winhofer, T. Scherer, M. Trauner, A. Kautzky-Willer, M. Krssak, M. Krebs, and P. Wolf. 2023. Increased GH/IGF-I Axis Activity Relates to Lower Hepatic Lipids and Phosphor Metabolism. J Clin Endocrinol Metab. 108:e989–e997. doi:10.1210/CLINEM/DGAD206.

Gierahn, T.M., M.H. Wadsworth, T.K. Hughes, B.D. Bryson, A. Butler, R. Satija, S. Fortune, J. Christopher Love, and A.K. Shalek. 2017. Seq-Well: portable, low-cost RNA sequencing of single cells at high throughput. Nature Methods 2017 14:4. 14:395–398. doi:10.1038/nmeth.4179.

Gomez Perdiguero, E., K. Klapproth, C. Schulz, K. Busch, E. Azzoni, L. Crozet, H. Garner, C. Trouillet, M.F. De Bruijn, F. Geissmann, and H.R. Rodewald. 2015. Tissue-resident macrophages originate from yolk-sac-derived erythro-myeloid progenitors. Nature. 518:547–551. doi:10.1038/nature13989.

Gong, T., C. Zhang, X. Ni, X. Li, J. Li, M. Liu, D. Zhan, X. Xia, L. Song, Q. Zhou, C. Ding, J. Qin, and Y. Wang. 2020. A time-resolved multi-omic atlas of the developing mouse liver. Genome Res. 30:263–275. doi:10.1101/GR.253328.119/-/DC1.

Hafemeister, C., and R. Satija. 2019. Normalization and variance stabilization of single-cell RNA-seq data using regularized negative binomial regression. Genome Biol. 20:1–15. doi:10.1186/S13059-019-1874-1/FIGURES/6.

Haynes, W. 2013. Wilcoxon Rank Sum Test. Encyclopedia of Systems Biology. 2354–2355. doi:10.1007/978-1-4419-9863-7_1185.

Heieis, G.A., T.A. Patente, L. Almeida, F. Vrieling, T. Tak, G. Perona-Wright, R.M. Maizels, R. Stienstra, and B. Everts. 2023. Metabolic heterogeneity of tissue-resident macrophages in homeostasis and during helminth infection. Nature Communications 2023 14:1. 14:1–15. doi:10.1038/s41467-023-41353-z.

Herzog, R., K. Schuhmann, D. Schwudke, J.L. Sampaio, S.R. Bornstein, M. Schroeder, and A. Shevchenko. 2012. LipidXplorer: a software for consensual cross-platform lipidomics. PLoS One. 7. doi:10.1371/JOURNAL.PONE.0029851.

Hiller, K., J. Hangebrauk, C. Jäger, J. Spura, K. Schreiber, and D. Schomburg. 2009. MetaboliteDetector: comprehensive analysis tool for targeted and nontargeted GC/MS based metabolome analysis. Anal Chem. 81:3429–3439. doi:10.1021/AC802689C.

Huang, H., I. Splichalova, N. Balzer, L. Seep, E.F. Taveras, C. Radwaniak, K. Mauel, N. Blank-Stein, N. Makdissi, J. Zurkovic, A. Kayvanjoo, K. Wunderling, M. Jessen, M. Yaghmour, T. Ulas, S. Grein, J. Schultze, Z. Liu, F. Ginhoux, M. Beyer, C. Thiele, J. Hasenauer, D. Wachten, and E. Mass. 2023. Developmental programming of Kupffer cells by maternal obesity causes fatty liver disease in the offspring. doi:10.21203/RS.3.RS-3242837/V1.

Hughes, T.K., M.H. Wadsworth, T.M. Gierahn, T. Do, D. Weiss, P.R. Andrade, F. Ma, B.J. de Andrade Silva, S. Shao, L.C. Tsoi, J. Ordovas-Montanes, J.E. Gudjonsson, R.L. Modlin, J.C. Love, and A.K. Shalek. 2020. Second-Strand Synthesis-Based Massively Parallel scRNA-Seq Reveals Cellular States and Molecular Features of Human Inflammatory Skin Pathologies. Immunity. 53:878–894.e7. doi:10.1016/J.IMMUNI.2020.09.015.

Humphrey, S.J., G. Yang, P. Yang, D.J. Fazakerley, J. Stöckli, J.Y. Yang, and D.E. James. 2013. Dynamic adipocyte phosphoproteome reveals that akt directly regulates mTORC2. Cell Metab. 17:1009–1020. doi:10.1016/j.cmet.2013.04.010.

Jacome-Galarza, C.E., G.I. Percin, J.T. Muller, E. Mass, T. Lazarov, J. Eitler, M. Rauner, V.K. Yadav, L. Crozet, M. Bohm, P.-L. Loyher, G. Karsenty, C. Waskow, and F. Geissmann. 2019. Developmental origin, functional maintenance and genetic rescue of osteoclasts. Nature. 568:541–545. doi:10.1038/s41586-019-1105-7.

Jin, S., C.F. Guerrero-Juarez, L. Zhang, I. Chang, R. Ramos, C.H. Kuan, P. Myung, M. V. Plikus, and Q. Nie. 2021. Inference and analysis of cell-cell communication using CellChat. Nature Communications 2021 12:1. 12:1–20. doi:10.1038/s41467-021-21246-9.

Kang, S., Y. Nakanishi, Y. Kioi, D. Okuzaki, T. Kimura, H. Takamatsu, S. Koyama, S. Nojima, M. Nishide, Y. Hayama, Y. Kinehara, Y. Kato, T. Nakatani, T. Shimogori, J. Takagi, T. Toyofuku, and A. Kumanogoh. 2018. Semaphorin 6D reverse signaling controls macrophage lipid metabolism and anti-inflammatory polarization. Nat Immunol. 19:561–570. doi:10.1038/S41590-018-0108-0.

Kayvanjoo, A.H., I. Splichalova, D.A. Bejarano, H. Huang, K. Mauel, N. Makdissi, D. Heider, H.M. Tew, N.R. Balzer, E. Greto, C. Osei-Sarpong, K. Baßler, J.L. Schultze, S. Uderhardt, E. Kiermaier, M. Beyer, A. Schlitzer, and E. Mass. 2024. Fetal liver macrophages contribute to the hematopoietic stem cell niche by controlling granulopoiesis. Elife. 13. doi:10.7554/ELIFE.86493.

Kineman, R.D., M. del Rio-Moreno, and A. Sarmento-Cabral. 2018. 40 YEARS of IGF1: Understanding the tissue-specific roles of IGF1/IGF1R in regulating metabolism using the Cre/loxP system. J Mol Endocrinol. 61:T187–T198. doi:10.1530/JME-18-0076.

Kineman, R.D., M. del Rio-Moreno, and D.J. Waxman. 2024. Liver-specific actions of GH and IGF1 that protect against MASLD. Nat Rev Endocrinol. doi:10.1038/S41574-024-01037-0.

Li, L., H. Zhou, J. Wang, J. Li, X. Lyu, W. Wang, C. Luo, H. Huang, D. Zhou, X. Chen, L. Xu, and P. Li. 2023. Metabolic switch from glycogen to lipid in the liver maintains glucose homeostasis in neonatal mice. J Lipid Res. 64. doi:10.1016/J.JLR.2023.100440.

Liu, Z., Y. Gu, S. Chakarov, C. Bleriot, I. Kwok, X. Chen, A. Shin, W. Huang, R.J. Dress, C.-A. Dutertre, A. Schlitzer, J. Chen, L.G. Ng, H. Wang, Z. Liu, B. Su, and F. Ginhoux. 2019. Fate Mapping via Ms4a3-Expression History Traces Monocyte-Derived Cells. Cell. 178:1509–1525.e19. doi:10.1016/j.cell.2019.08.009.

Luecken, M.D., and F.J. Theis. 2019. Current best practices in single-cell RNA-seq analysis: a tutorial. Mol Syst Biol. 15. doi:10.15252/MSB.20188746.

Mao, Q., L. Yang, L. Wang, S. Goodison, and Y. Sun. 2015. SimplePPT: A simple principal tree algorithm. Proc West Mark Ed Assoc Conf. 792–800. doi:10.1137/1.9781611974010.89.

Mass, E., I. Ballesteros, M. Farlik, F. Halbritter, P. Gunther, L. Crozet, C.E. Jacome-Galarza, K. Handler, J. Klughammer, Y. Kobayashi, E. Gomez-Perdiguero, J.L. Schultze, M. Beyer, C. Bock, and F. Geissmann. 2016. Specification of tissue-resident macrophages during organogenesis. Science (1979). 353:aaf4238–aaf4238. doi:10.1126/science.aaf4238.

Mass, E., and R. Gentek. 2021. Fetal-Derived Immune Cells at the Roots of Lifelong Pathophysiology. Front Cell Dev Biol. 9. doi:10.3389/fcell.2021.648313.

Mass, E., F. Nimmerjahn, K. Kierdorf, and A. Schlitzer. 2023. Tissue-specific macrophages: how they develop and choreograph tissue biology. Nature Reviews Immunology 2023. 1–17. doi:10.1038/s41577-023-00848-y.

Morgantini, C., J. Jager, X. Li, L. Levi, V. Azzimato, A. Sulen, E. Barreby, C. Xu, M. Tencerova, E. Näslund, C. Kumar, F. Verdeguer, S. Straniero, K. Hultenby, N.K. Björkström, E. Ellis, M. Rydén, C. Kutter, T. Hurrell, V.M. Lauschke, J. Boucher, A. Tomčala, G. Krejčová, A. Bajgar, and M. Aouadi. 2019. Liver macrophages regulate systemic metabolism through non-inflammatory factors. Nat Metab. 1:445–459. doi:10.1038/s42255-019-0044-9.

Müssig, K., H. Fiedler, H. Staiger, C. Weigert, R. Lehmann, E.D. Schleicher, and H.U. Häring. 2005. Insulin-induced stimulation of JNK and the PI 3-kinase/mTOR pathway leads to phosphorylation of serine 318 of IRS-1 in C2C12 myotubes. Biochem Biophys Res Commun. 335:819–825. doi:10.1016/J.BBRC.2005.07.154.

Nakagaki, B.N., K. Mafra, É. de Carvalho, M.E. Lopes, R. Carvalho-Gontijo, H.M. de Castro Oliveira, G. Henrique Campolina-Silva, C.D.M. de Miranda, M.M. Antunes, A.C.C. Silva, A.B. Diniz, D.M. Alvarenga, M.A. Freitas Lopes, V.A. de Souza Lacerda, M.S. Mattos, A.M. Araújo, P.V.T. Vidigal, C.X. Lima, G.A.B. Mahecha, M.F.M. Madeira, G.R. Fernandes, R.F. Nogueira, T.G. Moreira, B.A. David, R.M. Rezende, and G.B. Menezes. 2018. Immune and Metabolic Shifts During Neonatal Development Reprogram Liver Identity and Function. J Hepatol. doi:10.1016/J.JHEP.2018.08.018.

Nakanishi, Y., M. Izumi, H. Matsushita, Y. Koyama, D. Diez, H. Takamatsu, S. Koyama, M. Nishide, M. Naito, Y. Mizuno, Y. Yamaguchi, T. Mae, Y. Noda, K. Nakaya, S. Nojima, F. Sugihara, D. Okuzaki, M. Ikawa, S. Shimada, S. Kang, and A. Kumanogoh. 2024. Semaphorin 6D tunes amygdalar circuits for emotional, metabolic, and inflammatory outputs. Neuron. 112. doi:10.1016/J.NEURON.2024.06.017.

Nevado, C., A.M. Valverde, and M. Benito. 2006. Role of Insulin Receptor in the Regulation of Glucose Uptake in Neonatal Hepatocytes. Endocrinology. 147:3709–3718. doi:10.1210/EN.2005-1663.

Parker, B.L., G. Yang, S.J. Humphrey, R. Chaudhuri, X. Ma, S. Peterman, and D.E. James. 2015. Targeted phosphoproteomics of insulin signaling using data-independent acquisition mass spectrometry. Sci Signal. 8. doi:10.1126/SCISIGNAL.AAA3139/SUPPL_FILE/8_RS6_TABLES_S1_TO_S7.ZIP.

Pridans, C., A. Raper, G.M. Davis, J. Alves, K.A. Sauter, L. Lefevre, T. Regan, S. Meek, L. Sutherland, A.J. Thomson, S. Clohisey, S.J. Bush, R. Rojo, Z.M. Lisowski, R. Wallace, K. Grabert, K.R. Upton, Y.T. Tsai, D. Brown, L.B. Smith, K.M. Summers, N.A. Mabbott, P. Piccardo, M.T. Cheeseman, T. Burdon, and D.A. Hume. 2018. Pleiotropic Impacts of Macrophage and Microglial Deficiency on Development in Rats with Targeted Mutation of the Csf1r Locus. The Journal of Immunology. 201:2683–2699. doi:10.4049/JIMMUNOL.1701783.

Qiu, C., B.K. Martin, I.C. Welsh, R.M. Daza, T.M. Le, X. Huang, E.K. Nichols, M.L. Taylor, O. Fulton, D.R. O’Day, A.R. Gomes, S. Ilcisin, S. Srivatsan, X. Deng, C.M. Disteche, W.S. Noble, N. Hamazaki, C.B. Moens, D. Kimelman, J. Cao, A.F. Schier, M. Spielmann, S.A. Murray, C. Trapnell, and J. Shendure. 2024. A single-cell time-lapse of mouse prenatal development from gastrula to birth. Nature. 626:1084–1093. doi:10.1038/S41586-024-07069-W.

Revollo, J.R., A. Körner, K.F. Mills, A. Satoh, T. Wang, A. Garten, B. Dasgupta, Y. Sasaki, C. Wolberger, R.R. Townsend, J. Milbrandt, W. Kiess, and S. ichiro Imai. 2007. Nampt/PBEF/Visfatin Regulates Insulin Secretion in β Cells as a Systemic NAD Biosynthetic Enzyme. Cell Metab. 6:363–375. doi:10.1016/J.CMET.2007.09.003.

Rui, L., V. Aguirre, J.K. Kim, G.I. Shulman, A. Lee, A. Corbould, A. Dunaif, and M.F. White. 2001. Insulin/IGF-1 and TNF-alpha stimulate phosphorylation of IRS-1 at inhibitory Ser307 via distinct pathways. J Clin Invest. 107:181–189. doi:10.1172/JCI10934.

Rusin, D., L. Vahl Becirovic, G. Lyszczarz, M. Krueger, A. Benmamar-Badel, C. Vad Mathiesen, E. Sigurðardóttir Schiöth, K. Lykke Lambertsen, and A. Wlodarczyk. 2024. Microglia-Derived Insulin-like Growth Factor 1 Is Critical for Neurodevelopment. Cells. 13. doi:10.3390/CELLS13020184.

Saddi-Rosa, P., C.S. Oliveira, F.M. Giuffrida, and A.F. Reis. 2010. Visfatin, glucose metabolism and vascular disease: A review of evidence. Diabetol Metab Syndr. 2:1–6. doi:10.1186/1758-5996-2-21/TABLES/1.

Schultze, J.L., E. Mass, and A. Schlitzer. 2019. Emerging Principles in Myelopoiesis at Homeostasis and during Infection and Inflammation. Immunity. 50:288–301. doi:10.1016/j.immuni.2019.01.019.

Sud, M., E. Fahy, D. Cotter, K. Azam, I. Vadivelu, C. Burant, A. Edison, O. Fiehn, R. Higashi, K.S. Nair, S. Sumner, and S. Subramaniam. 2016. Metabolomics Workbench: An international repository for metabolomics data and metadata, metabolite standards, protocols, tutorials and training, and analysis tools. Nucleic Acids Res. 44:D463–D470. doi:10.1093/NAR/GKV1042.

Szklarczyk, D., A.L. Gable, D. Lyon, A. Junge, S. Wyder, J. Huerta-Cepas, M. Simonovic, N.T. Doncheva, J.H. Morris, P. Bork, L.J. Jensen, and C. Von Mering. 2019. STRING v11: protein-protein association networks with increased coverage, supporting functional discovery in genome-wide experimental datasets. Nucleic Acids Res. 47:D607–D613. doi:10.1093/NAR/GKY1131.

Traag, V.A., L. Waltman, and N.J. van Eck. 2019. From Louvain to Leiden: guaranteeing well-connected communities. Scientific Reports 2019 9:1. 9:1–12. doi:10.1038/s41598-019-41695-z.

Tye, L.M., and A.F. Burton. 1980. Glycogen deposition in fetal mouse tissues and the effect of dexamethasone. Biol Neonate. 38:265–269. doi:10.1159/000241375.

Viola, M.F., E. Franco Taveras, and E. Mass. 2024. Developmental programming of tissue-resident macrophages. Front Immunol. 15. doi:10.3389/FIMMU.2024.1475369.

Waraky, A., E. Aleem, and O. Larsson. 2016. Downregulation of IGF-1 receptor occurs after hepatic linage commitment during hepatocyte differentiation from human embryonic stem cells. Biochem Biophys Res Commun. 478:1575–1581. doi:10.1016/J.BBRC.2016.08.157.

Welz, L., N. Kakavand, X. Hang, G. Laue, G. Ito, M.G. Silva, C. Plattner, N. Mishra, F. Tengen, C. Ogris, M. Jesinghaus, F. Wottawa, P. Arnold, L. Kaikkonen, S. Stengel, F. Tran, S. Das, A. Kaser, Z. Trajanoski, R. Blumberg, C. Roecken, D. Saur, M. Tschurtschenthaler, S. Schreiber, P. Rosenstiel, and K. Aden. 2022. Epithelial X-Box Binding Protein 1 Coordinates Tumor Protein p53-Driven DNA Damage Responses and Suppression of Intestinal Carcinogenesis. Gastroenterology. 162:223–237.e11. doi:10.1053/J.GASTRO.2021.09.057.

Wolf, A., L. Moore, D. Lydahl, Ö. Naldemirci, M. Elam, and N. Britten. 2017. The realities of partnership in person-centred care: a qualitative interview study with patients and professionals. BMJ Open. 7. doi:10.1136/BMJOPEN-2017-016491.

Yan, X., E. Managlia, Y.Y. Zhao, X. Di Tan, and I.G. De Plaen. 2022. Macrophage-derived IGF-1 protects the neonatal intestine against necrotizing enterocolitis by promoting microvascular development. Communications Biology 2022 5:1. 5:1–13. doi:10.1038/s42003-022-03252-9.

Yang, L., W.H. Wang, W.L. Qiu, Z. Guo, E. Bi, and C.R. Xu. 2017. A single-cell transcriptomic analysis reveals precise pathways and regulatory mechanisms underlying hepatoblast differentiation. Hepatology. 66:1387–1401. doi:10.1002/HEP.29353/SUPPINFO.

Yang, L., X. Wang, J.X. Zheng, Z.R. Xu, L.C. Li, Y.L. Xiong, B.C. Zhou, J. Gao, and C.R. Xu. 2023. Determination of key events in mouse hepatocyte maturation at the single-cell level. Dev Cell. 58:1996–2010.e6. doi:10.1016/j.devcel.2023.07.006.

Young, M.D., and S. Behjati. 2020. SoupX removes ambient RNA contamination from droplet-based single-cell RNA sequencing data. Gigascience. 9. doi:10.1093/GIGASCIENCE/GIAA151.

Yu, G., L.G. Wang, Y. Han, and Q.Y. He. 2012. clusterProfiler: an R package for comparing biological themes among gene clusters. OMICS. 16:284–287. doi:10.1089/OMI.2011.0118.

Yu, G., L.G. Wang, G.R. Yan, and Q.Y. He. 2015. DOSE: an R/Bioconductor package for disease ontology semantic and enrichment analysis. Bioinformatics. 31:608–609. doi:10.1093/BIOINFORMATICS/BTU684.

Zappia, L., and A. Oshlack. 2018. Clustering trees: a visualization for evaluating clusterings at multiple resolutions. Gigascience. 7. doi:10.1093/GIGASCIENCE/GIY083.

